# Shaping tiny worlds: Paternal microbiota manipulation influences offspring microbial colonization and development in a sex role-reversed pipefish

**DOI:** 10.1101/2024.10.09.617486

**Authors:** Kim-Sara Wagner, Frédéric Salasc, Silke-Mareike Marten, Olivia Roth

**Affiliations:** Marine Evolutionary Biology, Zoological Institute, Kiel University, 24118 Kiel, Germany; OSU Institut Pytheas, Aix-Marseille Université, 13009 Marseille, France

## Abstract

Microbes are acquired through vertical and environmental horizontal transmission. Vertical transmission is directly linked to reproductive success and entails early transmission, facilitating coexistence of host and microbes over generations. The multiple potentially interacting routes of vertical transmission are challenging to be disentangled in conventional sex-role species, as they are mostly intermingled on the maternal side, i.e., through egg production, pregnancy, birth and postnatal care. The evolution of male pregnancy in syngnathids (pipefishes and seahorses) offers an opportunity to separate vertical microbial provisioning through the egg (maternal) from provisioning through pregnancy (paternal). We experimentally evaluated the existence and role of paternal microbiota provisioning through male pregnancy on offspring development and microbial colonization. Male pipefish were exposed to antibiotics, and subsequently recolonized with bacteria of paternal, maternal, and environmental origin (spike treatment). After pregnancy, the microbiome (16s rRNA) of developing offspring was repeatedly ribotyped. Paternal antibiotic and spike treatments influenced the microbial composition of the brood pouch and offspring microbiome development. Paternal spike treatment gave offspring a kickstart in life by reducing pregnancy duration and inducing offspring survival. Expanding on how distinct vertical microbial transmission routes shape offspring microbiome will foster our understanding of the holobiont function in health and disease.

## Introduction

The interplay of the host with its microbiome can be adaptive; facilitating long-lasting interactions that shape host development, digestion, immune regulation, behavior, and pathogen protection (He et al. 2020; Donald and Finlay 2023; Motta and Moran 2024). To understand the interactions of the host with its microbiome and their functional consequences on host development, we need to follow the microbes beginning at the initiation of their colonization. Vertical transmission fosters the initial microbial colonization as early as during egg development (Nyholm 2020), and can continue through the transfer in intimate parent-offspring interactions, such as pregnancy and postnatal parental care (Wosnick et al. 2022; Bogaert et al. 2023; Vitikainen et al. 2023). Vertical transmission routes are manifold and mostly bound to the maternal side impeding the disentangling of their roles in offspring microbial colonization and development. Therefore, the intertwining of egg production and pregnancy within the female body confounds the investigation of vertical transfer (Tanger et al. 2024), leaving us behind with open questions about the significance of the specific microbes transferred during egg production, pregnancy or postnatal parental care.

Throughout vertebrate evolution, viviparity — the development of the embryo inside the parent’s body — has evolved independently in over 150 lineages (Blackburn 2015). While in almost all cases the female is the pregnant sex, the syngnathids, a group of teleost fishes, represent the unique evolution of male pregnancy (Whittington and Friesen 2020). In the broad-nosed pipefish, *Syngnathus typhle*, the female deposits her eggs into the semi-enclosed brood pouch of the male where they are fertilized, protected, and nourished until the male gives birth (Stölting and Wilson 2007). This characteristic reproductive strategy provides the opportunity to disentangle the typically intermingled vertical microbial transmission through eggs and pregnancy: Both, transovarial microbial transfer by mothers and transfer during pregnancy by fathers may be equally expected (Supplementary Figure 1) (Tanger et al. 2024). In contrast, horizontal transfer was suggested to occur only when the brood pouch becomes strained and permeable during late pregnancy (Beemelmanns et al. 2019; Tanger et al. 2024) allowing to independently study the timing and role of vertical microbial transfer in depth.

The microbiome of *Syngnathus typhle* differs in composition between the female gonads and the male brood pouch (Beemelmanns et al. 2019). While the maternal microbiota seems to shape the offspring gut microbiome, paternal microbes rather influence the establishment of the offspring whole-body microbiome (Tanger et al. 2024). We aimed to provide the first experimental assessment of paternal microbial transmission through male pregnancy, by manipulating the paternal brood-pouch microbiome and disseminating its impacts on offspring microbiome and health. We depleted the paternal natural brood-pouch microbiome with antibiotics, and subsequently recolonized the brood pouch with selected natural bacterial isolates before mating males with untreated females. We hypothesized that this paternal microbial manipulation would influence offspring microbiome establishment and offspring development. Giving the intricate interplay of father and offspring during pregnancy, we expected paternal-specific microbes to successfully colonize the male tissue and be transferred to offspring. This study enhances the functional understanding of host-microbiota interactions in pregnancy and provides insights into the transmission and establishment of microbiota across generations.

## Methods

To ultimately assess the transfer of microbes through paternal pregnancy, we conducted three experiments: **(1)** Cultivation and characterization of sex-specific microbiota, **(2)** effectiveness of antibiotics for depleting the natural pipefish brood-pouch microbiome, and **(3)** manipulating paternal sex-specific microbes to unravel their impact on the offspring. In all three experiments, we sequenced the 16S rRNA gene (Bouchet et al. 2008) with Sanger sequencing (experiment 1) and with Illumina MiSeq Amplicon (V3/V4 region) (experiment 2 & 3). Data processing, visualization and statistical testing were performed in R Studio (R Core Team 2024).

### Fish catching and rearing

Broad-nosed pipefish (*Syngnathus typhle*) were caught in the bay of Orth (Fehmarn, Germany) (54°26’ N, 11°02’E). At the Helmholtz Centre for Ocean Research Kiel (GEOMAR), they were separated by sex, treated with formalin (1:8000 solution) on three consecutive days to remove potential ectoparasites, and kept in Baltic Sea water flow-through systems until allocation into the experimental facilities (16: 8 day-night; 15 PSU).

### (1) Cultivation and characterization of sex-specific microbiota

We aimed to cultivate the sex-specific microbiota of *Syngnathus typhle* from female eggs, male brood pouches throughout pregnancy and environmental samples to compare findings to a cultivation-independent NGS study (Beemelmanns et al. 2019). In total, 65 fish were used: 20 females, 17 non-pregnant, 16 early pregnant and 12 late pregnant males. Animals were euthanized with tricaine methane sulfonate (MS-222, 500 mg l^-1^; Sigma-Aldrich, St. Louis, USA) followed by weight and length measurements. Microbiota samples were isolated from six groups: female gonads (FG), non-pregnant males: placenta-like tissue of the developing brood pouch (NP), early pregnant males: placenta-like tissue (EPP) and whole juveniles (EPJ) as well as late pregnant males: placenta-like tissue (LPP) and whole juveniles (LPJ). Tissue samples were homogenized in autoclaved phosphate buffered saline (PBS) and plated onto Marine Broth (MB, Carl Roth GmbH, Karlsruhe, Germany) or *Vibrio*-selective TCBS agar plates (Carl Roth GmbH). After incubation, bacterial colonies were prepared for amplification of the 16S rRNA (primers 27F and 1492R). The PCR product was purified and Sanger sequenced (Wendling et al. 2017). Raw sequences were aligned and edited with CodonCode Aligner (V8.0.2). Consensus sequences were exported and submitted to NCBI’s BLAST for taxonomic identification within the 16S ribosomal RNA sequences (Megablast algorithm) (Altschul et al. 1997). The identified bacteria were compared to the sex-specific indicator species from Beemelmanns et al. (2019). Differences between stages and tissues were analyzed using PERMANOVA and ANOSIM on a ranked dissimilarity matrix and calculated with the Jaccard similarity coefficient (999 permutations). A multiple correspondence analysis (MCA), followed by pairwise Adonis analyzed microbial clusters of sexes, respectively pregnancy stages.

### (2) Effectiveness of antibiotics for natural pipefish microbiota depletion Microbiota sample collection

We assessed the effectivity of antibiotics in depleting the natural sex-specific microbiomes. We used 64 *Syngnathus typhle* (32 males, 32 females) and exposed them to antibiotics for 312 hours (13 days), consisting of 48 hours treatment, followed by 274 hours maintenance (reduction of antibiotic concentration by 75%). Pipefish were treated with chloramphenicol (treatment: 40 mg/L, maintenance: 10 mg/L), kanamycin (treatment: 10 mg/L, maintenance: 2.5 mg/L), a mixture of both antibiotics with identical concentrations (Munro et al. 1995; Verner-Jeffreys et al. 2003) or they were left untreated (control). Gut and sex-specific microbiota were sampled at 0, 6, 12, 24, 48, 192 and 312 hours. Gut microbiomes were sampled by gastric swabs using a sterile paper point (ANTÆOS® Absorbent Paper Points sterile, size 15/20/25, VDW GmbH). For assessing the reproductive sex-specific microbiomes, the inside of the brood pouch (male) and the surface of the ovipositor (female) were swabbed with sterile cotton swabs (Copan Italia, Thermo Fisher Scientific). Tank water antibiotic concentrations were assessed with agar well diffusion tests on *Epibacterium mobile* lawns (prevalent in pipefish, susceptible to both antibiotics applied) (Balouiri et al. 2016). After 48 and 312 four pipefish per treatment were euthanized (MS-222) and sampled for final gastric as well as reproductive sex-specific microbiota swabs. DNA was extracted using DNeasy 96 Blood & Tissue Kit (QIAGEN GmbH, Hilden, Germany) with enzymatic lysis buffer pre-treatment. The 16S rRNA V3/V4 was sequenced over Illumina Miseq amplicon sequencing at the Institute of Clinical Molecular Biology (IKMB, Kiel, Germany).

### Data analysis and statistics

From demultiplexed paired-end fastq files, sequences were merged, quality filtered, and analyzed using QIIME2 version 2019.10 (Caporaso et al. 2010). Chimeras were removed using the “consensus” method (DADA2, Callahan et al. 2016). Sequences were truncated 50 bases from forward and 70 bases from reverse reads in both runs. For taxa comparison, relative abundances based on all obtained reads were used. The Naïve Bayes classifier was trained on the SILVA138 99% OTUs full-length sequence database (Quast et al. 2013) and sequences from our dataset were assigned to ASVs from SILVA database with similarity on the 99% level. The phyloseq package v.1.46.0 (McMurdie and Holmes 2013) was used for further analyses and data visualization. The data set was subdivided for the reproductive sex-specific (brood pouch and ovipositor) and gut microbiomes. Count data was normalized, log-transformed and prevalence filtered to only preserve ASVs that are present in >20% of the samples. Alpha diversity indices were estimated for Shannon. Indicator species analysis using MULTIPATT with 999 permutations was performed using the R package indicspecies (Cáceres and Legendre 2009). PCAs and PCA biplots were calculated and visualized with FactoMineR (Lê et al. 2008). A mixed ANOVA (factors: treatment x time) tested for differences in Shannon diversity between treatment groups over time points. A repeated measured PERMANOVA evaluated the impact of treatments on sex-specific microbiota composition (beta diversity).

### (3) Manipulating paternal sex-specific microbiota to unravel impact on the offspring

In the last experiment, the methods established in the two previous experiments were applied: male pipefish were fully reciprocally treated with antibiotics (chloramphenicol) and recolonized with candidate strains for vertical and horizontal transfer (spike treatment) (Figure 1). We aimed to assess the potentially interacting impact of paternal antibiotic and spike treatment, on microbial colonization and offspring development.

**Figure 1:**
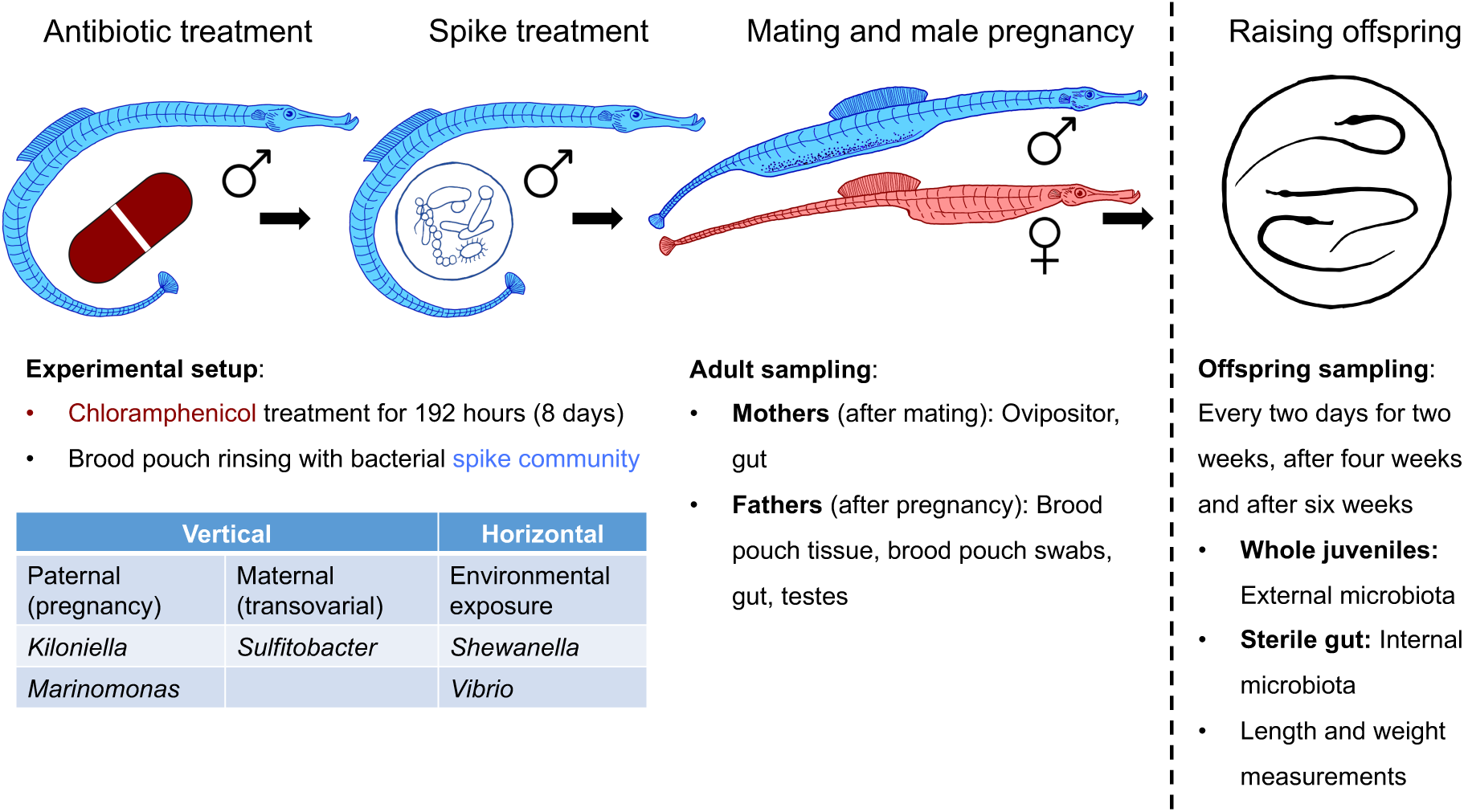
Setup of experiment 3: Manipulating paternal sex-specific microbiota to assess the impact on offspring development and early microbial colonization. Fathers received a treatment (antibiotics and/or spike) and were mated with untreated females. Offspring were kept in separate tanks and were repeatedly sampled for 45 days.

### Antibiotic treatment of male fish

72 *S. typhle* (36 males, 36 females) were used in this experiment. Half of the male fish were exposed to chloramphenicol in a high dose (40 mg/L) for 48 hours, followed by a low dose (10 mg/L) for 144 hours (6 days) (details: experiment 2). Females and untreated males were kept in separate circulation systems. Water temperatures were slowly increased from 11°C to 17°C. Fish were fed with live *mysid spp*., mysids for the antibiotic-treated males were treated with chloramphenicol (40 mg/L) for 1 hour prior to feeding.

### Spike treatment of male pipefish, pregnancy, and microbiota sampling of adults

Prior to mating, half of the male pipefish received a spike-bacteria treatment resulting in four different treatment groups: Control, Antibiotics (antibiotics applied, no spike bacteria added), Spike (no antibiotics applied, spike bacteria added) and AntiSpike (antibiotics and spike treatment). The spike community consisted of five selected bacterial strains cultivated during experiment 1: *Sulfitobacter* (vertical maternal transfer candidates), *Kiloniella* and *Marinomonas* (vertical paternal transfer candidates), *Shewanella* (horizontal transfer candidate) and *Vibrio* (prevalent in *S. typhle*). Overnight cultures were separately diluted to OD600 0.5, washed with phosphate-buffered saline (PBS), mixed equally and 30x concentrated (Supplementary Figure 7). Brood pouches of half of the males were flushed with 50µl spike culture. The other half was flushed with 50µl PBS (sham control). After treatment, pipefish pairs were put into 40 L glass mating tanks. After mating, females were removed and samples taken for gut and ovipositor microbiome ribotyping. After two weeks of pregnancy, the male fish were moved into individual 20 L tanks. Immediately after birth, placenta-like tissue, brood pouch surface, hind gut, and testes were sampled from the male pipefish. Tissues were flash frozen and stored at −80°C.

### Juvenile time series

10 juveniles from each batch were moved to 400 ml beakers for daily monitoring of length and weight for 45 days (Figure 1). From the remaining offspring (kept in their birth tanks), microbiota was sampled directly after birth, and then every two days for two weeks, after four weeks (28 days) and six weeks (45 days). Tissues were stored at −80°C. The fish were fed daily with live artemia. At each sampling, two offspring were sacrificed (one for whole-body and one for gut ribotyping), as well as food and seawater of each tank were collected. DNA was extracted and sequenced from all tissue and water samples for 16S rRNA ribotyping as in experiment 2. We only discuss relevant time points (complete dataset: supplementary spreadsheets 6 and 7).

### Data analysis

Paternal treatment effects on gestation length and number of offspring were assessed with a one-way ANOVA. Influence of paternal treatment on juvenile survival, weight and size development were tested in an ANCOVA with treatment as fixed factor and time as covariable, followed by pairwise comparisons and contrasts analyses. Processing of raw 16S rRNA amplicon sequences using the QIIME2 workflow followed the procedures described above. Truncation lengths were: run 63: forward 250, reverse 195; run 64: forward 260, reverse 190; run 65: forward 275, reverse 220; run 79: forward 250, reverse 195. Whole microbiome composition was assessed with a PCA biplot and MANOVA, followed by ANOVA. To identify sequences stemming from the recolonization spike culture, QIIME2 sequences were aligned to the sequences of the spike community. Relative abundance of spike community strains within the microbiome were analyzed with a generalized least square model (GLS) using normalized, log-transformed counts. Differences between treatment groups were assessed with a Kruskal-Wallis test followed by a Dunn’s post-hoc test.

## Results

### The sex-specific microbiota: Cultivation vs. Sequencing

We aimed to compare the isolated, cultivated and characterized sex-specific microbiota 16S rRNA Sanger sequences of *Syngnathus typhle* to the NGS dataset from Beemelmanns et al. (2019) facilitating the selection of strains suitable for manipulating the paternal sex-specific microbiota (experiment 3). 767 full length 16S rRNA sequences were trimmed (clip ends) and assembled, resulting in 376 consensus sequences (presence/absence matrix, species and genus level). From 79 tissue samples (13 female gonads (FG), 16 non-pregnant (NP), 11 early pregnancy pouch (EPP), 10 early pregnancy juveniles (EPJ), 12 late pregnancy pouch (LPP), 12 late pregnancy juveniles (LPJ)), 92 strains from 38 genera were identified. This corresponds to approximately 2.97% of the total number of 3090 OTUs (97% cut-off) identified in the NGS approach by Beemelmanns et al. (2019). Another six bacterial isolates from four genera were isolated from the seawater. Microbiome composition differed between stages (ANOSIM (p = 0.011); PERMANOVA (p = 0.001)): FG – LPJ: p = 0.020; NP – LPJ: p = 0.015; EPP – LPJ: p = 0.015; EPJ – LPJ: p = 0.023 (pairwise Adonis, Supplementary Table 2).

The MCA visualized associations between sexes or developmental stages with the 20 mainly contributing bacterial isolates (Supplementary Figure 3). Dimension 1 drove the microbiota of the female gonads (FG) away from the remaining five developmental stages, whereas dimension 2 segregated the microbiota associated to the late pregnancy stage (LPP, LPJ) from the other stages. Microbiota associated with non-pregnant (NP) males and the early pregnancy stage (EPP, EPJ) showed the strongest overlap. This pattern resembles the pattern identified by (Beemelmanns et al. 2019). *Vibrio* predominantly distinguished female eggs from other tissues, while *Pseudoalteromonas* was overrepresented in male brood pouches, particularly towards late pregnancy (Supplementary Figure 3B). In the NGS study, female eggs were represented by *Brevinema* and *Halomonas*, whereas the male brood pouch was characterized by *Marinomonas*, *Kiloniella*, *Ruegeria* and *Pseudoalteromonas (Beemelmanns et al. 2019)* (Complete dataset: supplementary spreadsheet 1).

For manipulating the sex-specific microbiota of *S. typhle* through recolonization, we pooled five bacterial strains with a potential role in transgenerational transfer and early microbial colonization (hereafter called “spike community”). As a similar pattern was identified in both studies but, not entirely driven by the same strains, we selected indicator strains from the NGS study (Beemelmanns et al. 2019) that were represented within the 2.97% cultivable part of the sex-specific pipefish microbiota. For paternal vertical transfer, we chose *Kiloniella*, *Marinomonas* and *Pseudoalteronomas*, indicators for the male brood pouch microbiome from non-pregnant to late pregnancy. For maternal vertical transfer, we selected *Sulfitobacter*, a strain present in female gonads and the brood pouch during late pregnancy but absent in non-or early pregnant males. We also included *Shewanella*, ubiquitous in aquatic ecosystems (Lemaire et al. 2020) isolated from water samples and in pipefish highly prevalent *Vibrio sinaloensis*.

### Impact of antibiotics on microbial diversity and on target strains for parental transfer

This experiment aimed to investigate how antibiotic treatments influenced sex-specific microbial diversity of *Syngnathus typhle* (males: brood pouch, females: ovipositor surface, pooled samples). 754 samples were submitted to 16S rRNA MiSeq amplicon sequencing. A total of 19,837,424 reads were returned (mean run 47: 28,517 reads/sample, mean run 48: 21,022 reads/sample), along with 10,688 representative sequences (Supplementary Table 1). The duration of the antibiotic treatment (time) as well as time x antibiotic interaction influenced sex-specific microbiota alpha diversity (Shannon) (ANOVA Type III: treatment: p = 0.629; time: p <.001; time x treatment: p <.001). No significant difference in alpha diversity were identified for timepoints T0 (p = 0.828), T6 (p = 0.165), T12 (p = 0.82) and T48 (p = 0.156) whereas from T24 (p = 0.002) onwards, diversity indices decreased (T192: p <.001; T312: p <.001). Within the first 48 hours (treatment), alpha diversity fluctuated (Figure 2A). After 8 days (maintenance), diversity was lowest in the treatments Chloramphenicol and Mix. After 13 days Chloramphenicol and Mix microbial diversities increased slightly but remained lower than diversities of Control and Kanamycin treatments, which remained high throughout the experiment (Figure 2A). This outcome corresponds to the well diffusion test results. Plates with water containing chloramphenicol (Chloramphenicol and Mix) showed clear zones of inhibition, whereas they were non-existent for in the other treatment groups (Kanamycin and Control) (Supplementary Figure 3). The alpha diversity of the gut microbiota was accordingly (Mixed ANOVA Type III on Shannon diversity: treatment: p = 0.276; time: p <.001; time x treatment: p <.001) (Supplementary Figure 4), which underlines the effectiveness of our chosen treatment (Complete dataset: supplementary spreadsheet 2).

**Figure 2:**
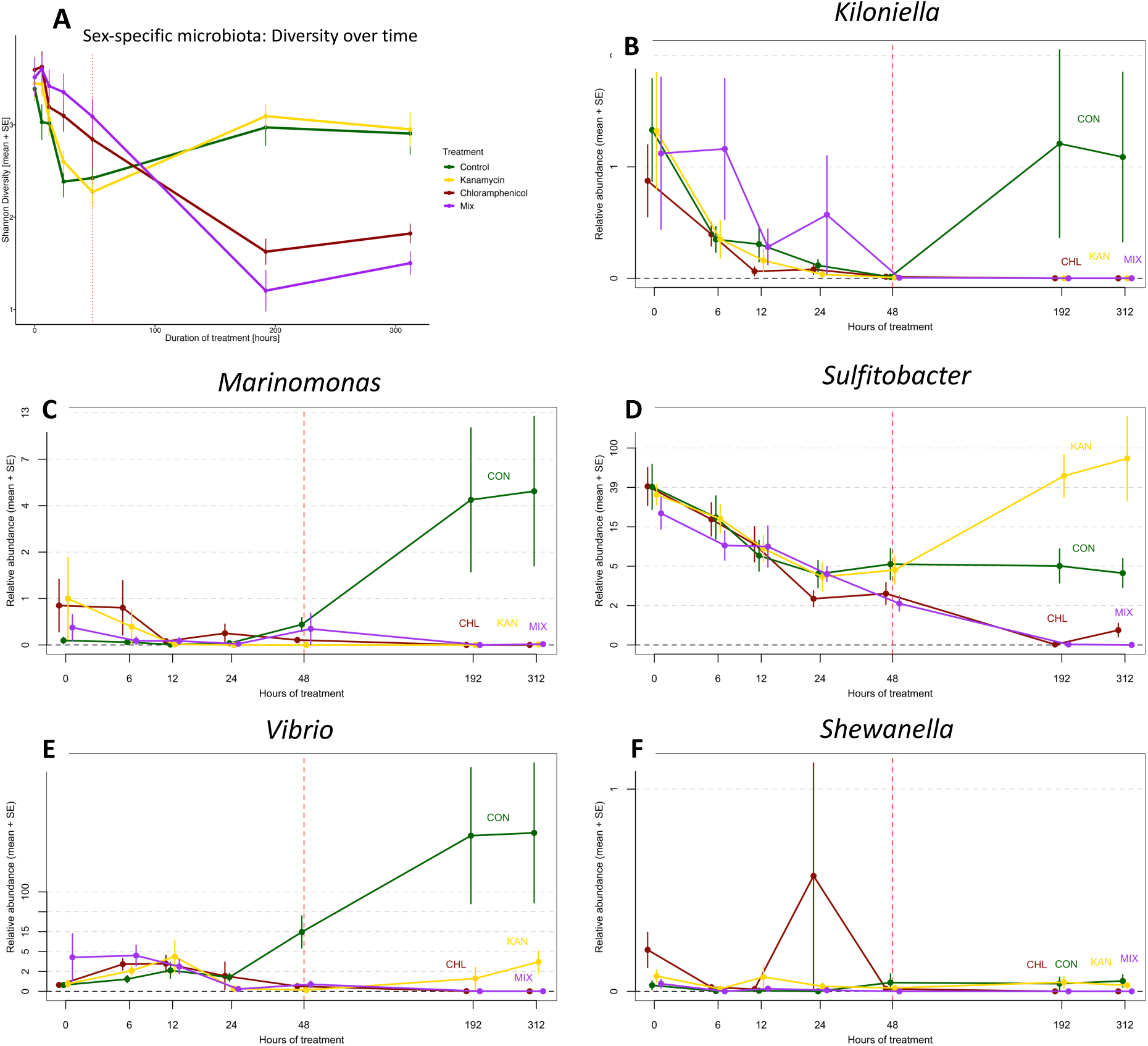
Impact of antibiotics (mean + SE) on Shannon diversity of sex-specific microbiota (A) and on relative abundance of spike community strains (B-F) over a period of 312 hours. Colours indicate treatment groups: Control (green), Kanamycin (yellow), Chloramphenicol (red) and Mix (purple). Red vertical line represents time of treatment dose (48 hours after treatment start).

To select spike community strains for manipulating the paternal sex-specific microbiota (experiment 3), we validated their successfully depletion by the antibiotic treatments. Five bacterial strains (Paternal: *Kiloniella*, *Marinomonas*; Maternal: *Sulfitobacter*; Environmental/ubiquitous: *Shewanella*, *Vibrio)* were depletable by chloramphenicol and could not be detected after 192 hours (Figure 2B-F). *Pseudoalteromonas* could not be depleted by chloramphenicol treatment and was therefore excluded from the spike community (Supplementary Figure 5).

### Impact of antibiotics on microbial community structure (beta diversity)

Similar to alpha-diversity, a repeated measures PERMANOVA suggested significant differences in treatment, time and the treatment x time interaction (all p < 0.001) of the sex-specific microbiota composition (beta-diversity, Supplementary Table 3).

Principal component analyses (PCA) of four time points (treatment: 6, 48 hours; maintenance: 192, 312 hours) suggested a similar microbiota composition in all treatments at the onset of the antibiotic application (MANOVA: T6: p = 0.457, Figure 3A). After 48 hours microbial communities separated into Chloramphenicol and Mix treated individuals vs. Kanamycin treated and control individuals (T48: p = 4.577e-09, Figure 3B). This separation intensified over time and was strongest after 192 hours (T192 and T312: p < 2e-16, Figure 3C-D).

**Figure 3:**
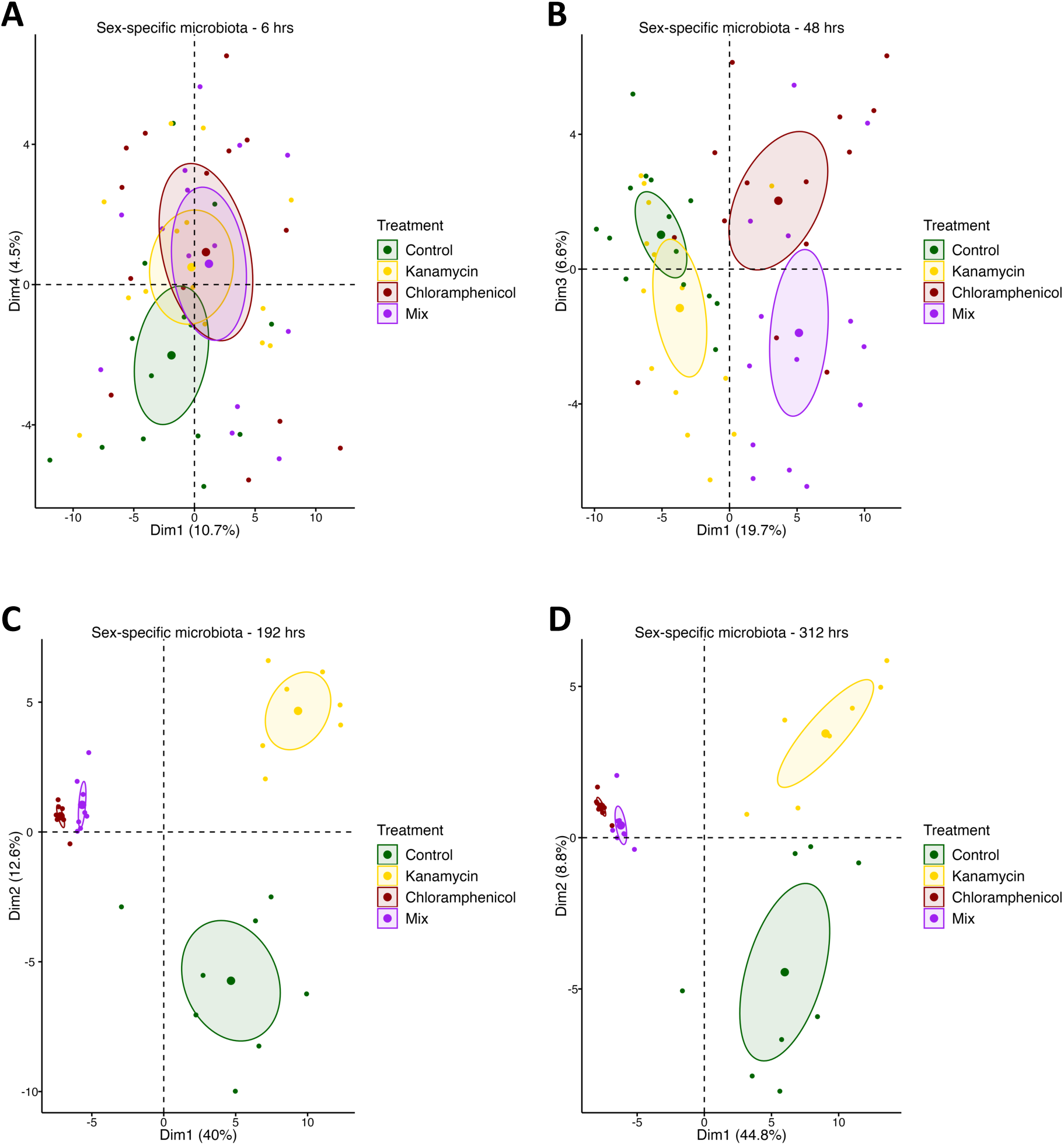
PCAs of antibiotic treatment (A: 6 hours, B: 48 hours, C: 192 hours, D: 312 hours) represent effectiveness of chloramphenicol. Normalized, log-transformed data include ASVs present in >20% of samples. Colours represent treatment group and ellipses 95% confidence intervals.

An indicator species analysis for T192 displayed associations between species patterns and combinations of groups, the 30 strains with the highest contribution to the observed patterns are displayed (Supplementary Figure 6). At T192, the Chloramphenicol-treated bacterial composition was driven by *Paraglaciecola* and *Maribacter*, whereas Kanamycin and Control-composition were driven by *Sulfitobacter* and *Pseudoalteromonas* (Complete dataset: supplementary spreadsheet 3).

The antibiotic application modified the bacterial composition and reduced its diversity. Chloramphenicol was suitable to deplete a large number of strains within the characterized microbiota and depleted the five isolates chosen as spike community for experiment 3. 192 hours antibiotic treatment appeared most suitable for experiment 3, minimizing stress and disease risk as opposed to longer treatments.

### Impact of antibiotic and spike treatments on pregnancy and parental tissues

Experiment 3 aimed to assess the impact of paternal antibiotic and spike treatment on pregnancy, early microbial colonization and offspring health from mating until 45 days post release (dpr). Fish were pregnant for 29.18 ± 1.62 days (mean + SD), spike treatment reduced the length of gestation (28.47± 0.8 days) (SP: p = 0.01). Proportion of brood pouch filled with eggs and number of offspring born was not influenced by treatments (Figure 4, Supplementary Table 4) (Complete dataset: supplementary spreadsheet 4).

**Figure 4:**
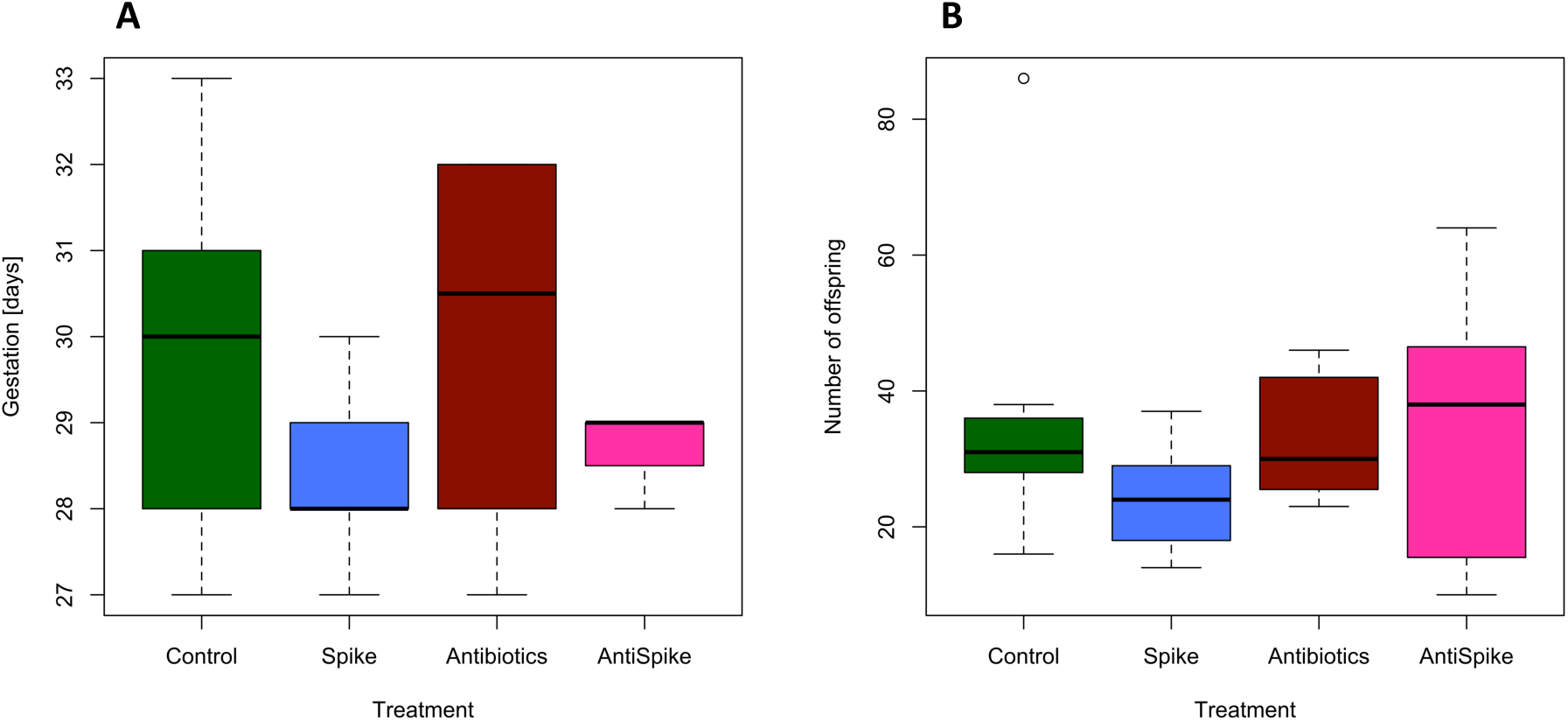
Gestation length (days) (A) is significantly shorter for males who received a spike treatment. In contrast, number of offspring (B) does not differ between males from different treatment groups

For microbiota ribotyping, a total of 1080 samples, including adult tissue, juvenile tissue and water samples, were 16S rRNA MiSeq amplicon sequenced. 34,021,625 reads were returned (mean run 63: 29,912 reads/sample, mean run 64: 27,566.5 reads/sample, mean run 65: 23,589 reads/sample, mean run 79: 135,136.5 reads/sample), along with 23,841 representative sequences (Supplementary Table 1). Male pipefish were sampled after giving birth (approximately day 30 post-treatment). We analyzed alpha diversity (Shannon) and beta diversity of the placenta-like tissue microbiome and its surface. Alpha diversity of the males was not influenced by treatment (Supplementary Figure 8, Supplementary Table 5). Antibiotic treated fish (Antibiotics and AntiSpike group) showed a significantly different microbial composition (beta diversity, p = 0.002) on the surface (brood pouch swabs) and within the placenta-like tissue (Figure 5A and 5C) compared to non-antibiotic treated fish (Control and Spike) (Figure 5A and 5C). Male gut microbiome was not influenced by the antibiotic treatment (Figure 5B). In contrast, the Spike group overlapped with the AntiSpike group and differed from the other groups (SP: p = 0.027). In the testes, all groups overlapped in their microbiome and no statistical differences were identified (Figure 5D). While males were sampled after giving birth, females (all untreated) were sampled directly after mating. Ribotyping the female gut and ovipositor microbiota allowed a) to illuminate sexual dimorphism in gut microbiome when comparing to gut microbiome of untreated males, and b) provided a positive control for the presence of our spike community, as the ovipositor of a mated female had been in direct contact with the spike-treated male brood pouch when mated with males from AntiSpike and Spike treatment groups (treatment assignment thus refers to the treatment of the male mating partner).

**Figure 5:**
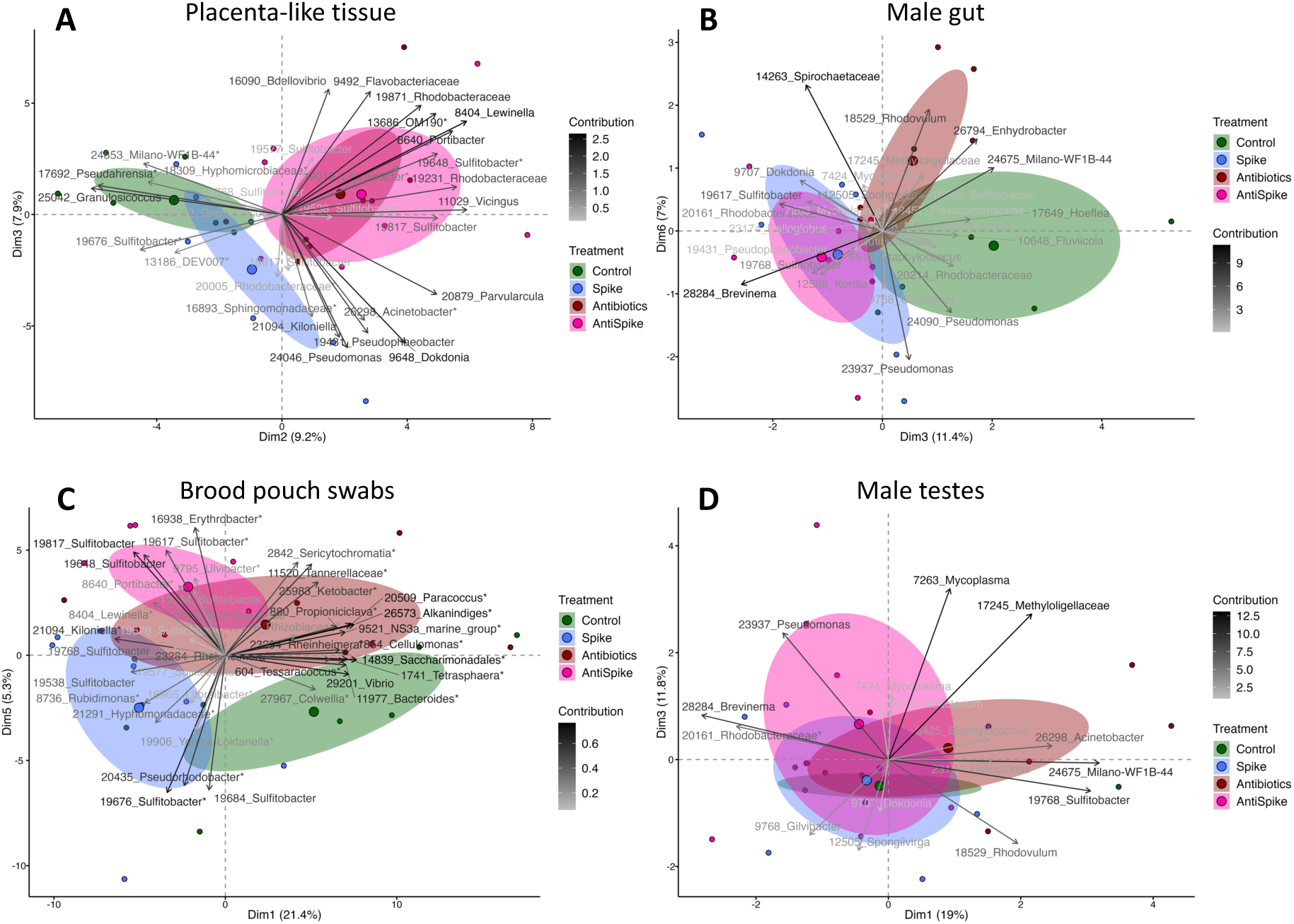
PCA biplots displaying treatment effects (antibiotics and spike) on paternal tissue. In the placenta-like tissue (A) and on the surface of the brood pouch (B), antibiotics affected the microbial composition. In the male gut (C) the spike treatment had a significant impact whereas in the testes (D), none of the treatments showed an effect. Colours represent the paternal treatment group: green = Control, blue = Spike, red = Antibiotics, pink = AntiSpike, Ellipses show a confidence interval around the group’s mean. Factor map shows ASV ID and genus name respectively family name.

### Treatment did not influence female pipefish microbiomes (Supplementary Figure 10)

Sex was an important factor explaining variation in alpha (p = 0.0002, Supplementary Figure 11, Supplementary Table 6) and beta diversity of gut microbial composition (p = 0.026, Supplementary Figure 12, Supplementary Table 7), suggesting a sexual microbiome dimorphism in untreated pipefish. In the gut tissue, this dimorphism was driven by the indicator species *Aliivibrio* in the female gut as well as *Rhodobacteriaceae*, *Maribacter*, *Enhydrobacter* and *Milano-WF1B-44* in the male gut (Supplementary Figure 12) (Complete dataset: supplementary spreadsheet 5).

### Impact of antibiotic and spike treatments on life-history and offspring microbiome

Offspring gained weight over time (time: p <2e-16) (Figure 6B, Supplementary Table 8), in contrast, offspring size was influenced by time and its interaction with treatment (time: p <2e-16, time*treatment: p = 0.045) (Figure 6C, Supplementary Table 9). Treatment, time and their interaction significantly affected offspring survival (treatment: p = 1.85e-14; time: p <2e-16; treatment*time: p = 8.03e-10) (Supplementary Table 10). Offspring of spike-treated fathers survived better than all other groups (Contrasts: Spike vs Antibiotics, AntiSpike and Control (all p < 0.0001, Figure 6A, Supplementary Table 11) (Complete dataset: supplementary spreadsheet 4).

**Figure 6:**
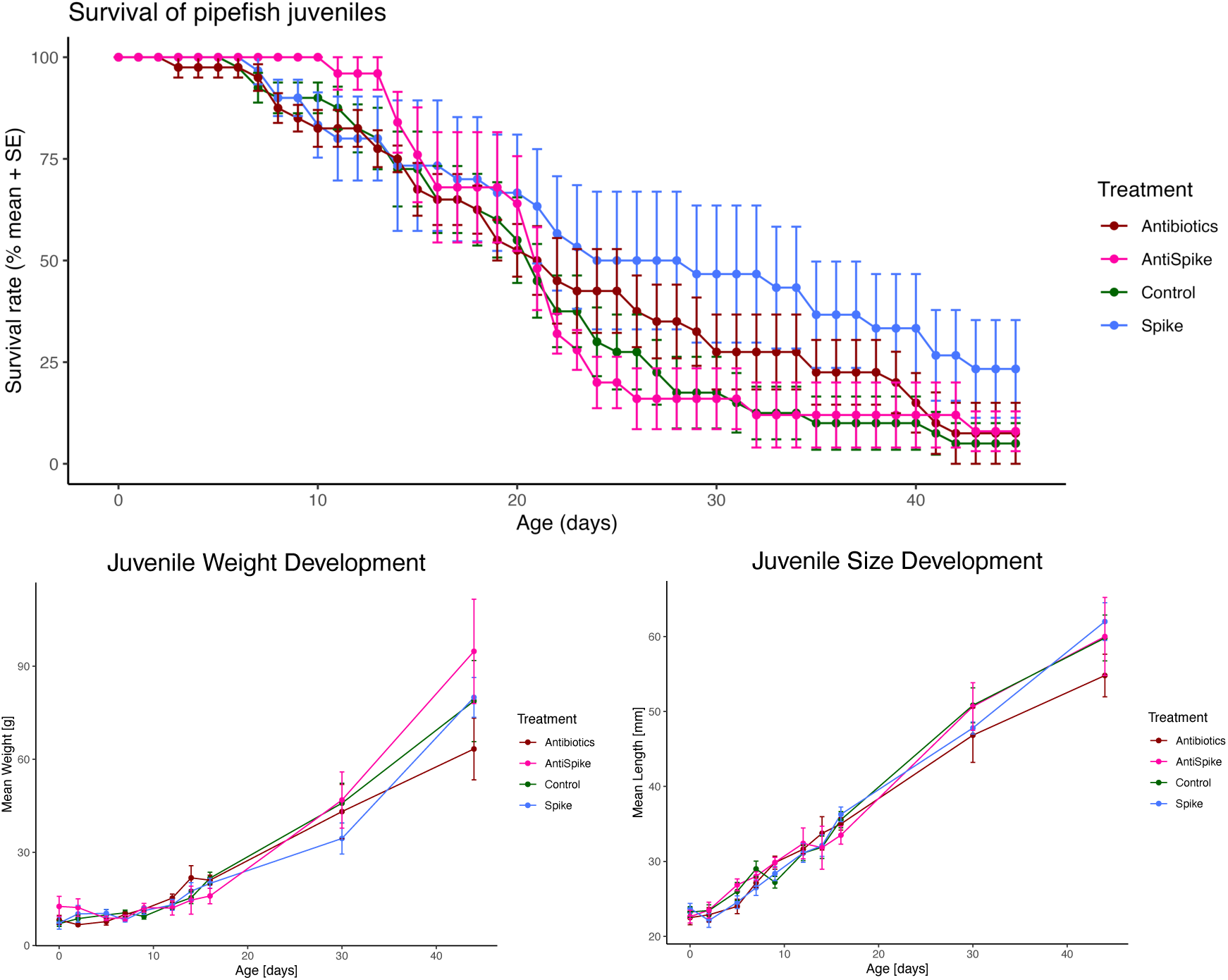
Life-history data of pipefish juveniles from birth until 45 days after birth. Survival rate (A), Weight development (B) and Size development (C). All plots show mean ± standard error (SE). Colour represents paternal treatment.

Upon offspring birth, juvenile whole-body and gut microbiomes were sequenced every second day for two weeks, after 30 and 45 days. Offspring from Control fathers had lower whole-body microbiota diversity across all time points (p = 0.039) (Alpha diversity, factor treatment), while the time point (age) itself had no significant impact (Figure 12A, Supplementary Table 12). Conversely, offspring gut microbiome changed (Alpha diversity) over time (p < 0.001), unaffected by the paternal treatment (Figure 12B, Supplementary Table 13). At birth, offspring whole-body microbiome was unaffected by paternal treatment (dpr 0). However, two days after birth, the spike treatment significantly shaped offspring microbiome (p = 0.013), while paternal antibiotics treatment indicated a trend (p = 0.052). Paternal treatment effects were strongest between six and ten days after birth (antibiotics: dpr 6: p = 0.039; dpr 10: p = 0.025) and (spike: dpr 6: p = 0.006; dpr 10: p = 0.017), but remained detectable up to 30 days after birth (antibiotics: p = 0.025; spike: p = 0.009) (Supplementary Table 14, Figure 7). The absence of a treatment effect at dpr 8 and dpr 12 indicates fluctuations in paternal treatment effects.

**Figure 7:**
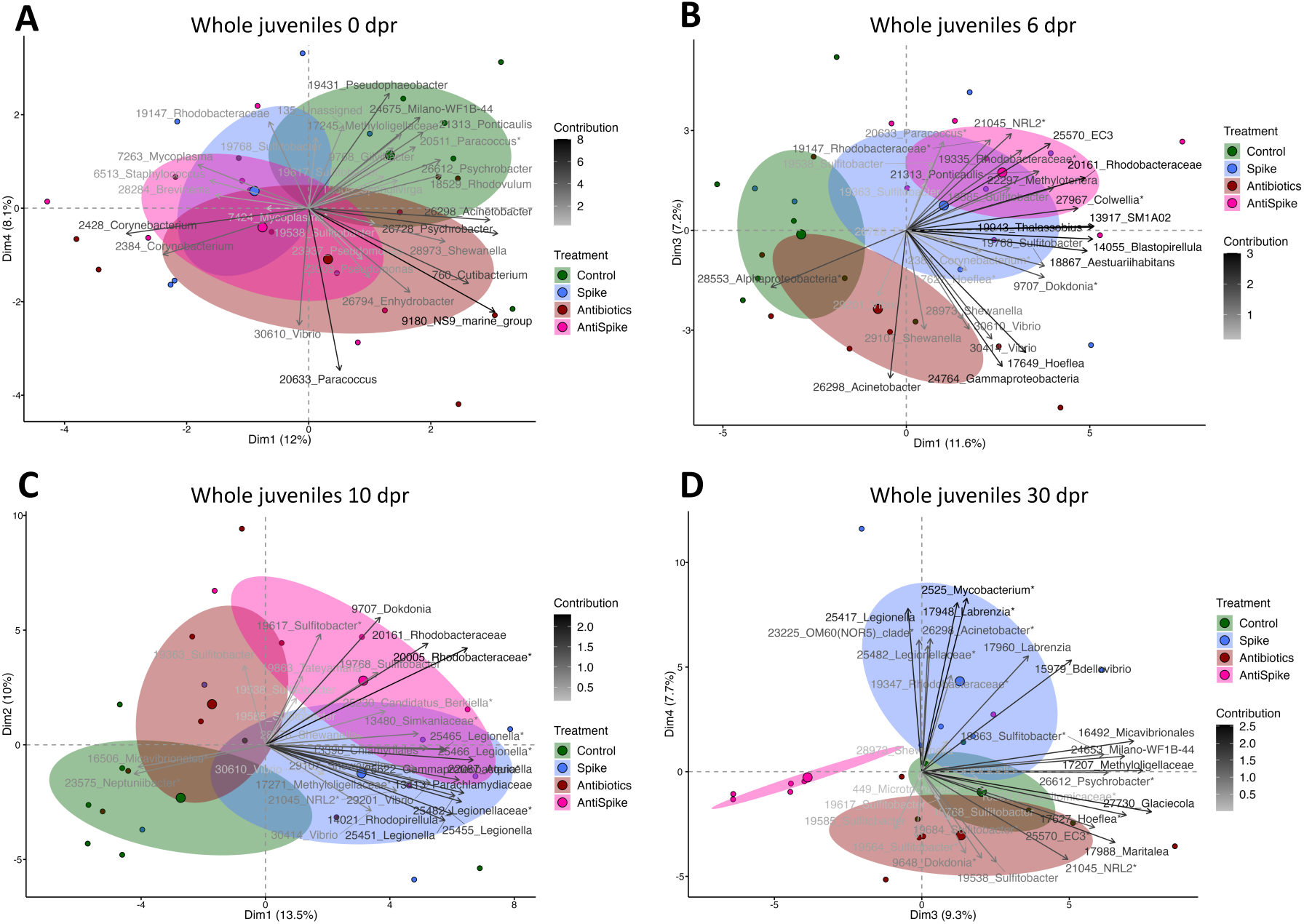
Shifts in offspring whole body microbiome composition from birth (A), 6 days (B), 10 days (C) up to 30 days (D) after birth. Colours represent the paternal treatment group. Ellipses show confidence intervals. Factor map shows ASV ID and genus name respectively family name.

Paternal spike treatment did not shape offspring gut microbiome, with the exception of 12 days post-release (p = 0.044; Supplementary Table 15, Supplementary Figure 13) (Complete dataset: supplementary spreadsheet 6 and 7).

### Searching for the spike bacteria in the adults and offspring

When examining the 16S rRNA sequences of our spike community in the relevant tissues after approximately two (females) respectively 30 days of pregnancy (males), we found identical sequence hits across all tissues (both adults and juveniles) and treatment groups, underlying that the microbes used in our spike community are members of the natural pipefish microbiome. In tissue of spike-treated fathers (Spike and AntiSpike), relative abundances of spike community members were undistinguishable from untreated fathers (Control and Antibiotics) (Figure 8A, Supplementary Figure 14). Normalized counts of Spike members were higher on the surface of male brood pouches (brood pouch swabs) from spike treated fathers (p = 0.044), a pattern driven by high *Shewanella* prevalence (p = 0.018). However, across all adult tissues and treatments *Shewanella* counts were high, while *Marinomonas* was mostly absent indicating a lack of colonization (Supplementary Table 16).

**Figure 8:**
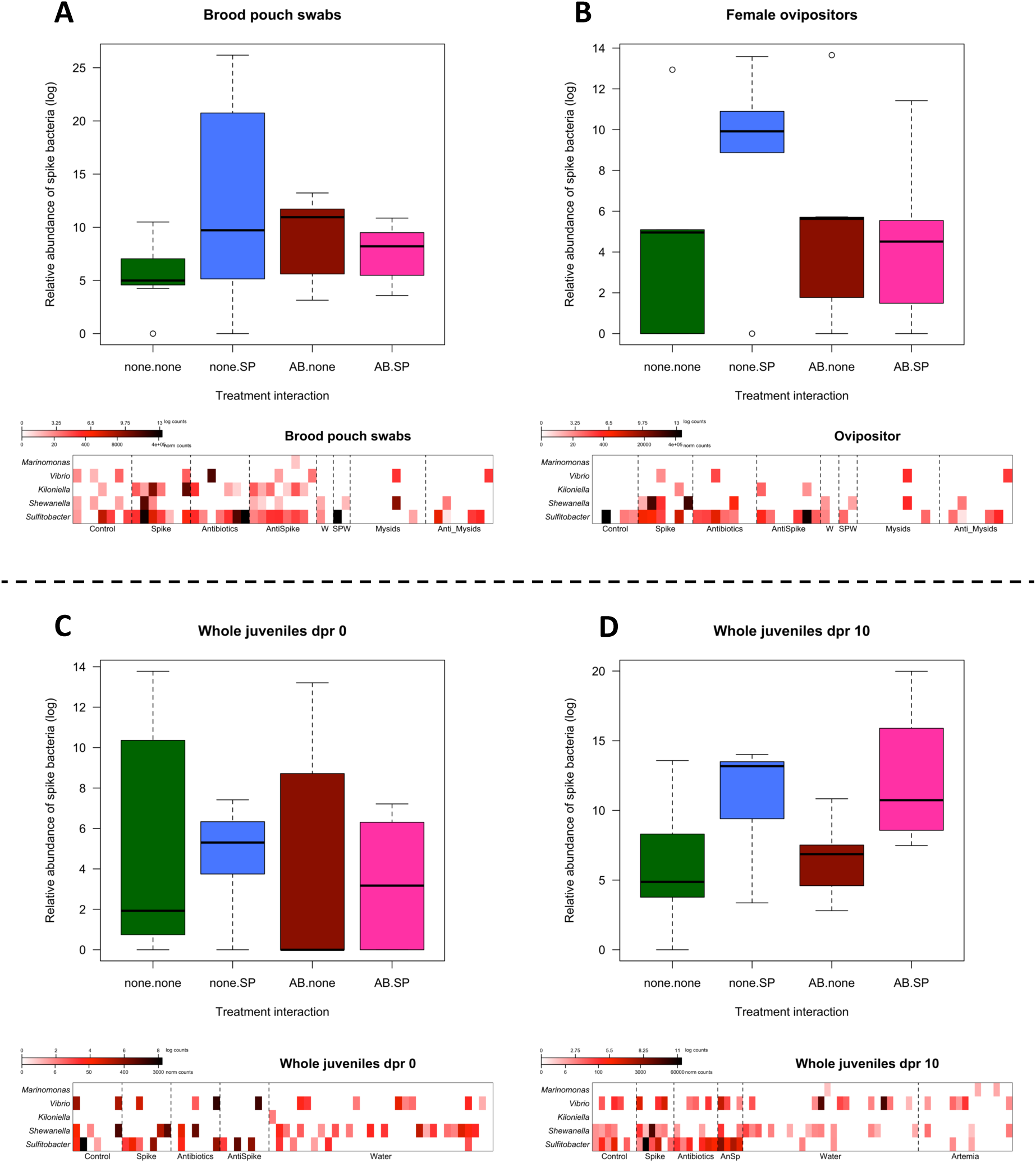
Prevalence of spike bacteria. Upper boxplots show relative abundance of spike bacteria within the whole microbiome of the respective tissue. Colours represent paternal treatment group. Lower heatmaps show individual counts (normalized) per sample. Darker colour indicates higher counts. A: Swabs of brood pouch surface, B: Female ovipositor after contact with treated male brood pouch, C: Whole body of juveniles after birth, D: whole body of juveniles 10 days after birth.

**Figure 9:**
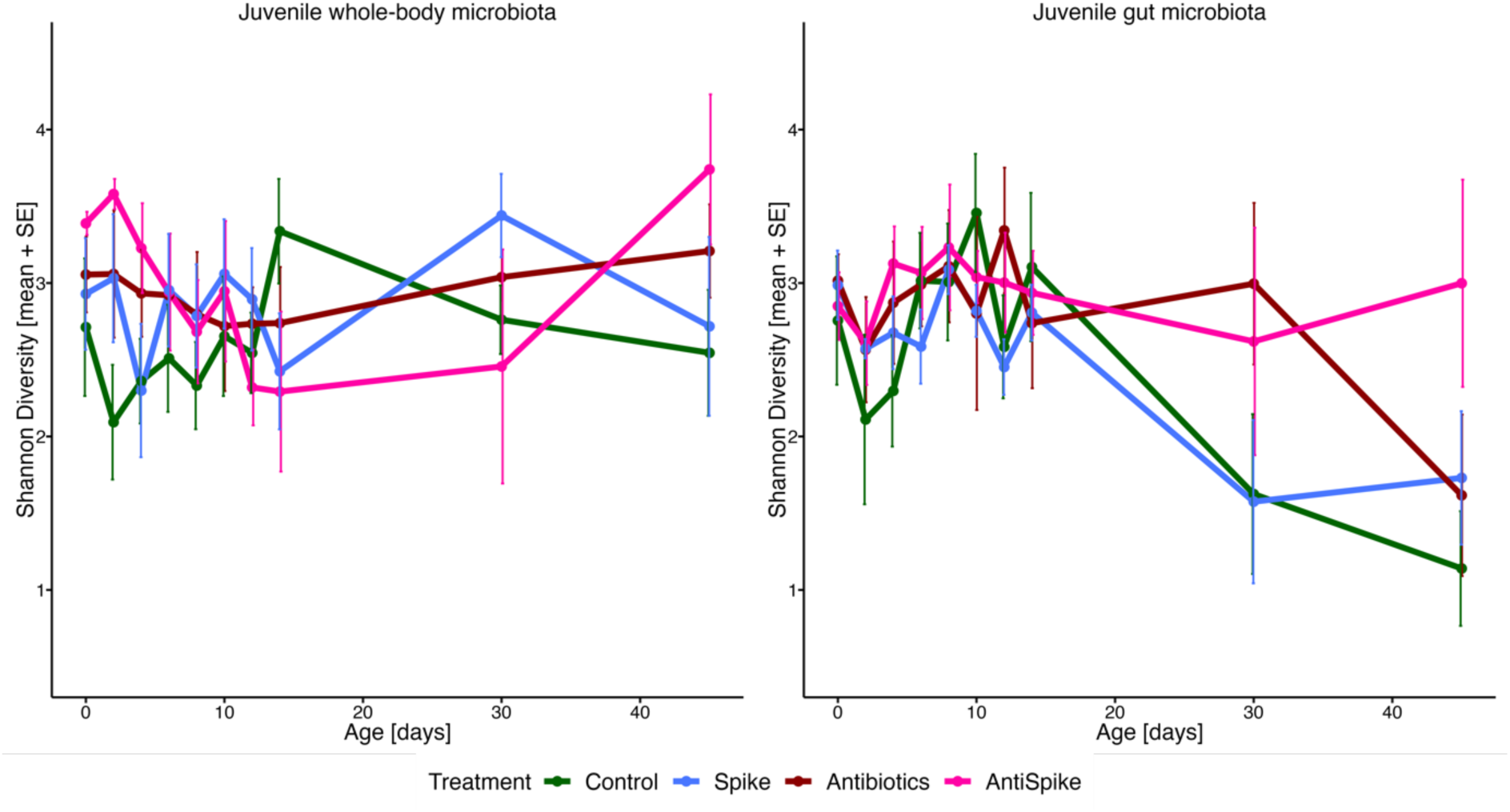
Alpha diversity (Shannon index) of juvenile whole body (A) and gut (B) microbiota from birth until 45 days post release. Colors indicate paternal treatment groups. Data points represent mean diversity ± standard error (SE).

Paternal spike treatment (spike and AntiSpike) induced spike community prevalence in offspring whole-body microbiome at 10 days after birth (p = 0.013), driven by high counts of *Shewanella* (p = 0.018), particularly in the AntiSpike treatment group (Figure 8D). No similar tendency was identified for offspring gut microbiome. *Kiloniella* was absent from juvenile gut samples, suggesting a lack of transfer respectively colonization. Regardless of paternal treatment, *Marinomonas* was absent in the majority of offspring samples (both whole-body-microbiome and gut-microbiome), but prevalent in the microbiome of the food (artemia, Figure 8D). Since this strain did not colonize adult tissue (Supplementary Table 16), the rare counts for this strain in our offspring dataset probably stem from food. *Vibrio* counts in the offspring should be treated with caution, as a ubiquitous strain in the natural environment of the pipefish, *Vibrio* was particularly abundant in artemia (Complete dataset: supplementary spreadsheet 8).

## Discussion

We aimed to experimentally assess the existence and role of paternal microbiota provisioning on offspring life-history and early microbial colonization through male pregnancy in the sex-role reversed pipefish *Syngnathus typhle*. To achieve this goal, we cultivated and characterized the sex-specific pipefish microbiota (experiment 1) and studied the effectiveness of antibiotics in depleting its microbiota (experiment 2). This facilitated to select microbes for paternal microbial manipulation (experiment 3) to unravel how paternal microbial provisioning influenced offspring development and microbiome establishment.

Both cultivation-dependent and cultivation-independent approaches (Beemelmanns et al. 2019) supported the presence of distinct microbial communities in female eggs, the brood pouch of non-pregnant males, and throughout male pregnancy. Early offspring microbial colonization was tissue-specific, shaped rather independently by mother and father (Tanger et al. 2024).

The here observed sexual dimorphism in microbial communities, coupled with a decreased beta diversity in females (Supplementary Figure 11, 12), strikingly contrasts with findings from conventional sex role species (de la Cuesta-Zuluaga et al. 2019; Valeri and Endres 2021). Sex roles and life history, rather than sex alone, seem to be the drivers of immunological activity and microbial diversity (Roth et al. 2010; Pappert et al. 2024). The differences in investment into parental care and secondary sexual ornaments appear to extend into the complex interactions between the microbiome and the immune system. Investigating parental microbial provisioning will help to understand how sex-role dynamics influence health and development across generations.

By treating pipefish with chloramphenicol, we successfully decreased pipefish brood pouch microbial diversity. Chloramphenicol was particularly effective against the microbes later assigned as experimental spike community, consisting of microbes from parental origin and isolates ubiquitous in pipefish and their natural environment. Achieving a completely germ-free status in our fish remained unattainable with antibiotic treatment alone. Working with *S. typhle*, a non-model marine species, presents challenges for gnotobiotic rearing due to its obligatory male pregnancy and the largely unknown impact of environmental microbes on its development and health. We thus acknowledge the inherent bias of persistent microbes, that we were unable to exclude throughout the conducted experiments.

Treating paternal pipefish with antibiotics influenced the composition and diversity of their microbiomes. The antibiotic treatment effect remained detectable 30 days post-treatment (after male pregnancy) when assessing the microbiome of the placenta-like tissue and its surface, but was absent when assessing the microbiome of the male gut or testes. This suggests that the antibiotic treatment affected the parts of the body, which were in direct contact with the water, and penetrated the underlying tissues, while it did not have a long-lasting effect on internal organs. After the antibiotic treatment, the gut microbiome might have reverted towards a stable core microbiome during the four weeks of pregnancy. In contrast, a long-lasting influence of the spike treatment on the microbiome composition was observed in the male gut despite that bacteria were added directly into the brood pouch. To shed light on time dynamics of potentially interacting antibiotic and spike treatments on parental tissues, ribotyping the microbial composition of paternal tissue throughout male pregnancy would be necessary. We have decided against this approach in the concurrent experiment to avoid disturbing or sacrificing the developing embryos due to limited housing opportunity in our experiment (36 breeding pairs in separate tanks, 9 per treatment).

Paternal spike treatment (Spike and AntiSpike) shortened paternal pregnancies compared to those of unexposed fathers (Control and Antibiotics). Also, in mammalian pregnancy various types of bacteria have been linked to preterm labor (Daskalakis et al. 2023). However, contrary to health challenges observed upon mammalian preterm birth, offspring from spike-treated fathers survived better than those from all other groups, suggesting spike treatment to be adaptive for offspring health.

After offspring left the paternal brood pouch and the associated protection from environmental microbial colonization, paternal spike as well as antibiotic treatment started to influence the whole-body microbiome up to 30 days after birth (Figure 7B-D, Supplementary Table 14). This suggests that the shift in the paternal brood pouch microbiome upon paternal treatment must have influenced the offspring microbial composition and the susceptibility of the offspring towards microbial colonization. The latter could explain why paternal treatment effects only emerged after birth, when offspring were suddenly exposed to a cocktail of surrounding environmental bacteria. In contrast, the offspring gut microbiome was not influenced by paternal treatment. While an impact on offspring gut microbiome by paternal treatment was potentially masked by a strong influence of the bacteria consumed over artemia feeding, this finding supports the notion that pipefish fathers primarily shape the whole-body microbiome, while mothers shape the gut microbiome (Tanger et al. 2024).

Despite initiating the spike community treatment with a mixture of pure bacterial cultures (Supplementary Figure 7), sequences matching our spike bacteria were found across all tissues (adults and juveniles) and treatment groups. This either suggests that the respective bacteria also colonized through environmental sources, making experimentally applied and natural strains difficult to detect. Or, non-mutually exclusive, that the sequences obtained from 16S V3/V4 ribotyping were too conserved to reliably detect the specific strains provided in the spike community. In particular *Shewanella* and *Vibrio* often exhibit high 16S rRNA copy numbers, leading to taxonomic classification differences for the same strain (Větrovský and Baldrian 2013). Metagenomics could have better distinguished bacterial strains, however the high sample size ribotyped throughout the experiment (n = 1080) rendered this impossible.

Although this experiment focused solely on paternal microbial transmission during pregnancy, the selected bacteria originated from paternal, maternal, and environmental sources, permitted to assess the microbial specificity of transgenerational effects. *Sulfitobacter*, identified as maternally-specific strain in experiment 1, exhibited high prevalence upon inoculation (Spike and Antispike) both in paternal tissue post-pregnancy and offspring tissues. Also, in offspring tissues of Control and Antibiotics treated fathers, it was present, albeit in lower counts. This may suggest that the extent of its colonization and transfer depended on the overall prevalence of the strain, especially given its high concentrations in the spike treatment, or that *Sulfitobacter* was simply ubiquitous in the environment and therefore horizontally colonized the tissues after antibiotic effects ceased. Conversely, *Marinomonas*, previously identified as paternal-specific in all studies on *S. typhle* microbiota, was found only in low abundances on the brood pouch surface within the AntiSpike group and was absent in all offspring tissues. Similarly, *Kiloniella*, which was present in high abundances in the paternal placenta-like tissue upon spike exposure, could not be detected in offspring. It is tempting to speculate that while these isolates are naturally sex-specific, they were here outcompeted by other isolates of maternal or environmental origin upon paternal inoculation.

In line with earlier studies, paternal spike treatment only shaped the whole-body microbiome (Tanger et al. 2024). This underlines the importance in disentangling routes of vertical bacteria transfer, as maternal transfer was suggested to rather shape offspring gut microbiomes (Tanger et al. 2024), giving both parents specific roles in microbial inheritance. In natural systems such as *S. typhle*, differentiating between environmental and parental effects on microbial colonization and development is essential. The interplay between these factors is complex and requires comprehensive analysis to fully understand the patterns involved. Due to the discussed limitations of our approach, future research should apply genetic-fluorescent- and antibiotic resistance-tagged spike strains. This would facilitate a detailed exploration of microbial transfer pathways from parents to offspring, providing precise insights into how specific bacteria are transmitted and established across generations, as well as pinpointing their functions in both paternal as well as offspring microbiomes.

By assessing the parental as well as offspring immune responses upon both antibiotic and spike exposures, we could gain deeper insights into the complex interactions between microbial communities, environmental factors, and host immune systems across generations, linking to previously suggested bi-parental trans-generational immune priming (Roth et al. 2012; Beemelmanns and Roth 2016, 2017). Paternal spike bacteria did not only shape offspring microbiomes but also increased offspring survival at a larger size. Insights into adaptive paternal effects on offspring microbiomes can add to our fundamental understanding of the ecological and evolutionary contexts of microbial interactions enlightening the adaptive significance of microbiota in host biology.

## Conclusion

Our study underscores the significant role of both paternal and environmental factors in shaping the microbiota of offspring. It also highlights the necessity for advanced methodologies and comprehensive studies to unravel the complexities of microbial transfer and its implications for host health. Continued research in this area will not only enhance our understanding of microbial ecology but may also contribute to managing microbial communities for promoting health and preventing disease.

This work highlights the role of paternal provisioning during male pregnancy, demonstrating that early colonization patterns in pipefish juveniles are primarily tissue-rather than sex-or bacterial strain-specific. This indicates that a dysbiosis in the father’s sex-specific tissues may notably influence offspring microbial colonization, as demonstrated here for the maternal indicator *Sulfitobacter*. While these findings might slip our attention in a balanced ecosystem, they could gain significance in the context of global change, potentially leading to the transfer of an altered microbial community with unknown impacts on offspring health and development. Our study examines only a small part of a complex and dynamic system, while the impact of environmental bacterial remains to be predicted in future studies. This underscores the need for advanced methodologies, such as fluorescent tagging, to better understand microbial transfer pathways in more detail. Future research should focus on exploring these transfer mechanisms and their long-term effects on offspring health to provide a more comprehensive understanding of the ecological and evolutionary implications of microbial inheritance.

## Authors’ contributions

**KSW**: Conceptualization, investigation, data curation, formal analysis, methodology, project administration, resources, software, validation, visualization, writing; **FS**: investigation, methodology, data curation, formal analysis, visualization; **SMM**: Conceptualization, methodology; investigation **OR**: conceptualization, funding acquisition, investigation, methodology, project administration, data analysis, resources, supervision, validation, writing. All authors gave final approval for publication and agreed to be held accountable for the work performed therein.

## Conflict of interest declaration

The authors declare to have no competing interests.

## Funding

This work was funded by the European Research Council (ERC) under the European Union’s Horizon research and innovation program (MALEPREG; eu-repo/grantAgreement/EC/H2020/755659) granted to O.R and supported by the Collaborative Research Center (CRC) 1182 “Origin and Function of Metaorganism” funded by the German Research Foundation.

## Acknowledgements

We express our gratitude to Katja Cloppenborg-Schmidt, Diana Gill, and Rebekka Leßke for their support in the laboratory and assistance with library preparation. We also thank Fabian Wendt, Johannes Hasse, and the student assistants for their dedicated care of the fish and maintenance of the aquaria facilities. Special thanks to Sören Franzenburg and the IKMB team for the Illumina MiSeq Sequencing. We are particularly thankful to Ralf Schneider, Arseny Dubin and Isabel Tanger for their exceptional contributions to our statistical analysis, data curation, and continuous guidance throughout the data analysis process. Finally, we are grateful to Thorsten Reusch, the Marine Evolutionary Biology group, and the members of the CRC1182 Origin and Function of Metaorganism for their support and inspiring discussions. Their collaboration created a highly stimulating environment for host-microbe interaction research at GEOMAR and Kiel University.

## Supplementary Material

**Supplementary Figure 1:**
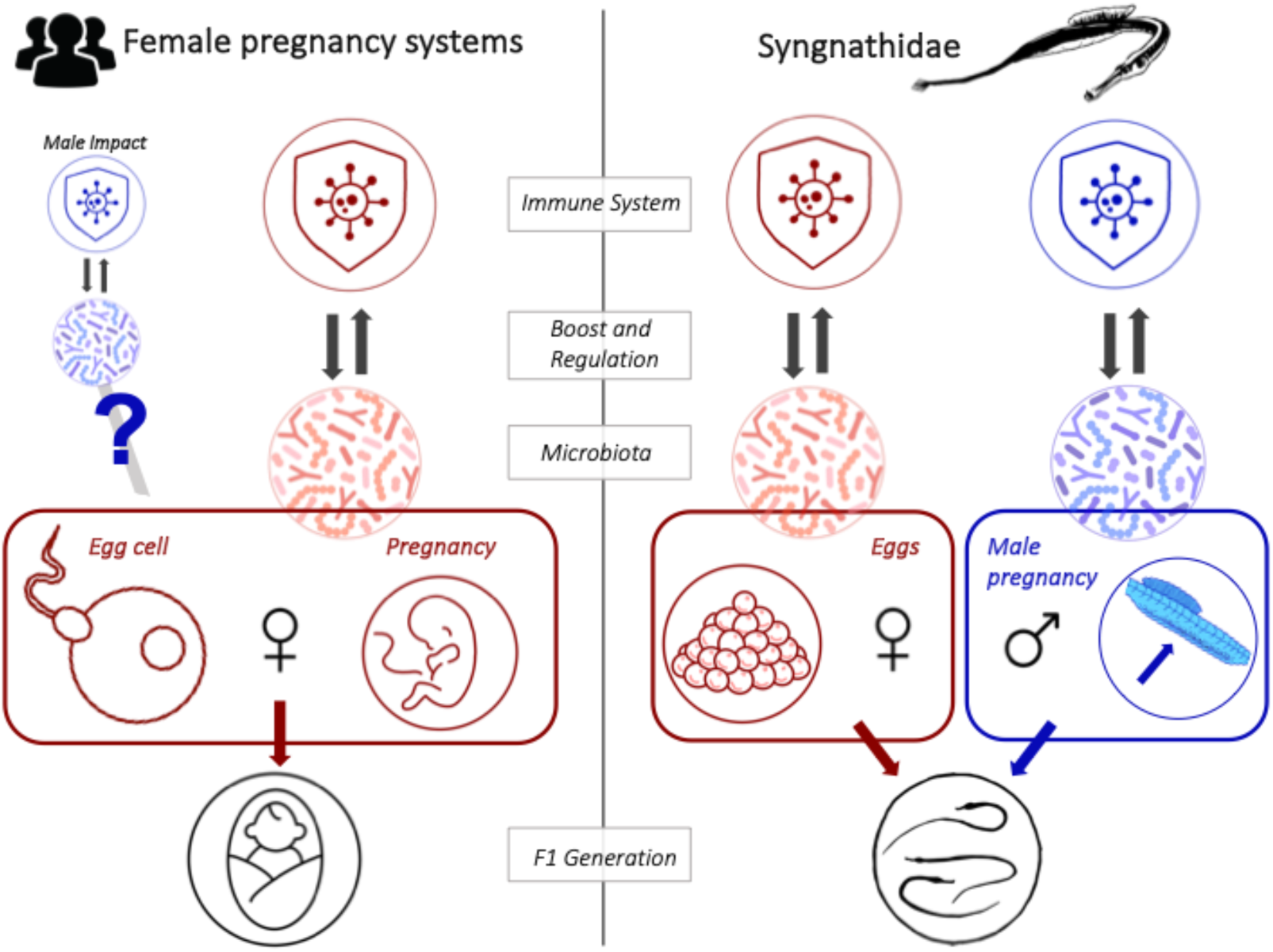
Immune system and microbiota transfer across generations: Comparison of classic female pregnancy systems (leV) with unique male pregnancy in the pipefish Syngnathus typhle (right) with a focus on the interaction of the microbiota and the immune system.

**Supplementary Table 1:** QIIME2 sequence processing statistics. Samples were randomly distributed over all sequencing runs. DADA2 workflow was applied separately for each run before merging all runs into a master data set.

| MiSeq Run | 47 | 48 | 63 | 64 | 65 | 79 |
| --- | --- | --- | --- | --- | --- | --- |
| Experiment | Experiment 2 |  | Experiment 3 |  |  |  |
| Samples | 377 | 377 | 289 | 384 | 377 | 30 |
| Min. reads | 22 | 22 | 13 | 7 | 16 | 57639 |
| Median | 28517.0 | 21022.0 | 29912.0 | 27566.5 | 23589.0 | 135136.5 |
| Mean | 30871.65 | 21747.52 | 30129.50 | 28620.82 | 26122.26 | 149190.37 |
| Max. reads | 102427 | 55732 | 83458 | 115401 | 137303 | 344546 |
| Total reads | 11638610 | 8198814 | 8707426 | 10990395 | 9848093 | 4475711 |
| Rep seqs | 10688 |  | 23841 |  |  |  |

<u>Experiment 1:</u> Cultivation and characterization of sex-specific microbiota

**Supplementary Table 2:** Pairwise adonis comparing sexes and/or stages of pregnancy in S. typhle. Abbreviations: FG = female gonads, NP = non pregnant, EPP = early pregnant pouch, EPJ = early pregnant juveniles, LPP = late pregnant pouch, LPJ = late pregnant juveniles.

| Pairs | Df | SumsOfSqs | F.Model | R2 | p.value | p.adjusted |
| --- | --- | --- | --- | --- | --- | --- |
| FG - NP | 1 | 0.6658655 | 1.4589270 | 0.05126430 | 0.035 | 0.1050000 |
| FG - EPP | 1 | 0.5352662 | 1.1910171 | 0.05367113 | 0.176 | 0.3016667 |
| FG - EPJ | 1 | 0.4758822 | 1.0656973 | 0.04829656 | 0.369 | 0.4612500 |
| FG - LPP | 1 | 0.5355834 | 1.1764959 | 0.04866279 | 0.167 | 0.3016667 |
| FG - LPJ | 1 | 0.7346852 | 1.6352001 | 0.06637657 | 0.004 | **0.0200000** |
| NP - EPP | 1 | 0.3875046 | 0.8660220 | 0.03482752 | 0.689 | 0.7382143 |
| NP - EPJ | 1 | 0.5451620 | 1.2252517 | 0.04857243 | 0.181 | 0.3016667 |
| NP - LPP | 1 | 0.5849061 | 1.2918955 | 0.04733623 | 0.087 | 0.2175000 |
| NP - LPJ | 1 | 0.9318041 | 2.0822734 | 0.07414903 | 0.001 | **0.0150000** |
| EPP - EPJ | 1 | 0.3080231 | 0.7114310 | 0.03802120 | 0.891 | 0.8910000 |
| EPP - LPP | 1 | 0.4249862 | 0.9564965 | 0.04564200 | 0.540 | 0.6230769 |
| EPP - LPJ | 1 | 0.8262708 | 1.8886978 | 0.08628644 | 0.002 | **0.0150000** |
| EPJ - LPP | 1 | 0.4849914 | 1.0990119 | 0.05208831 | 0.328 | 0.4612500 |
| EPJ - LPJ | 1 | 0.7400999 | 1.7034783 | 0.07848872 | 0.006 | **0.0225000** |
| LPP - LPJ | 1 | 0.4841525 | 1.0888442 | 0.04715889 | 0.354 | 0.4612500 |

**Supplementary Figure 2:**
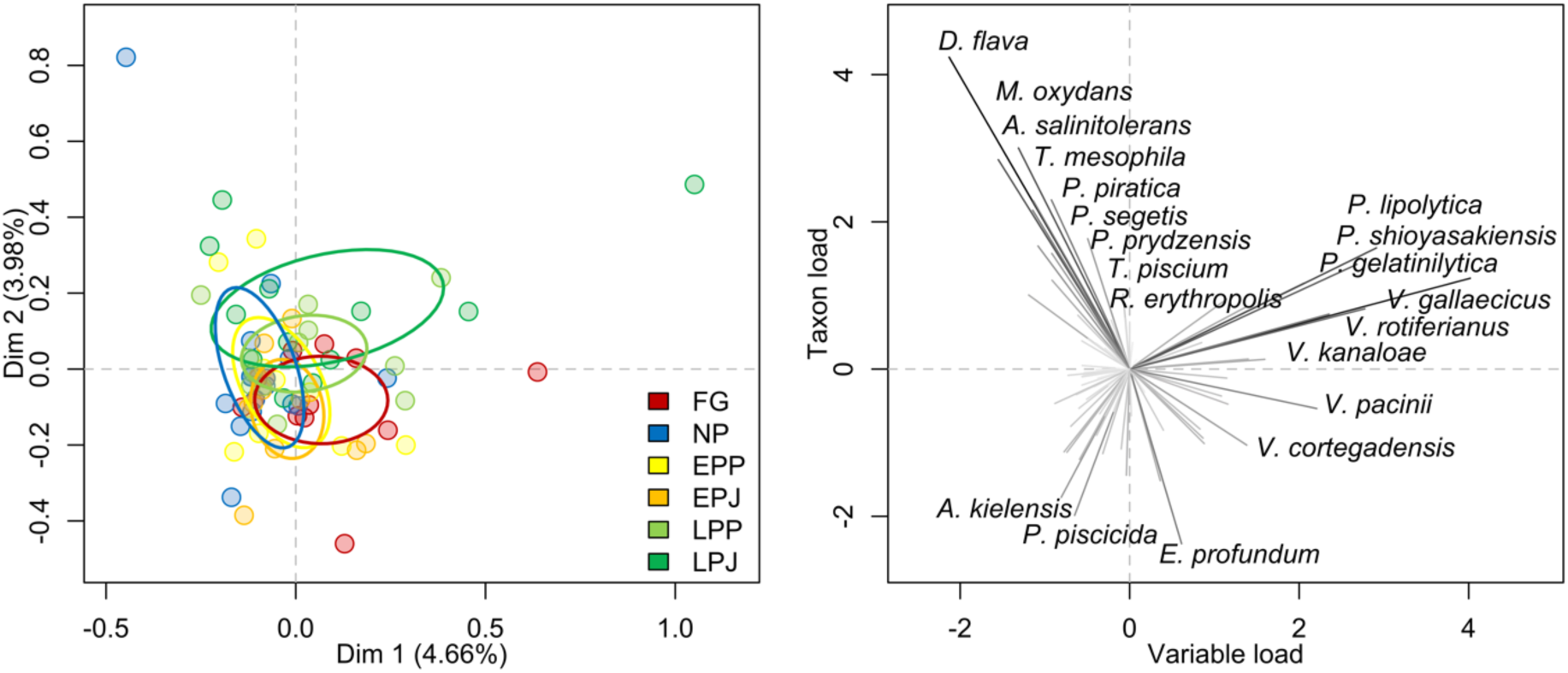
Multiple correspondence analysis (MCA) of six groups for dimensions 1 and 2 (A) and factor map displaying the loadings retained by the isolated bacterial species (B). Ellipses include 40% of data of the respective group. Colors represent sex and pregnancy stages as described above.

<u>Experiment 2</u>: Effectiveness of antibiotics for natural pipefish microbiota depletion

**Supplementary Figure 3:**
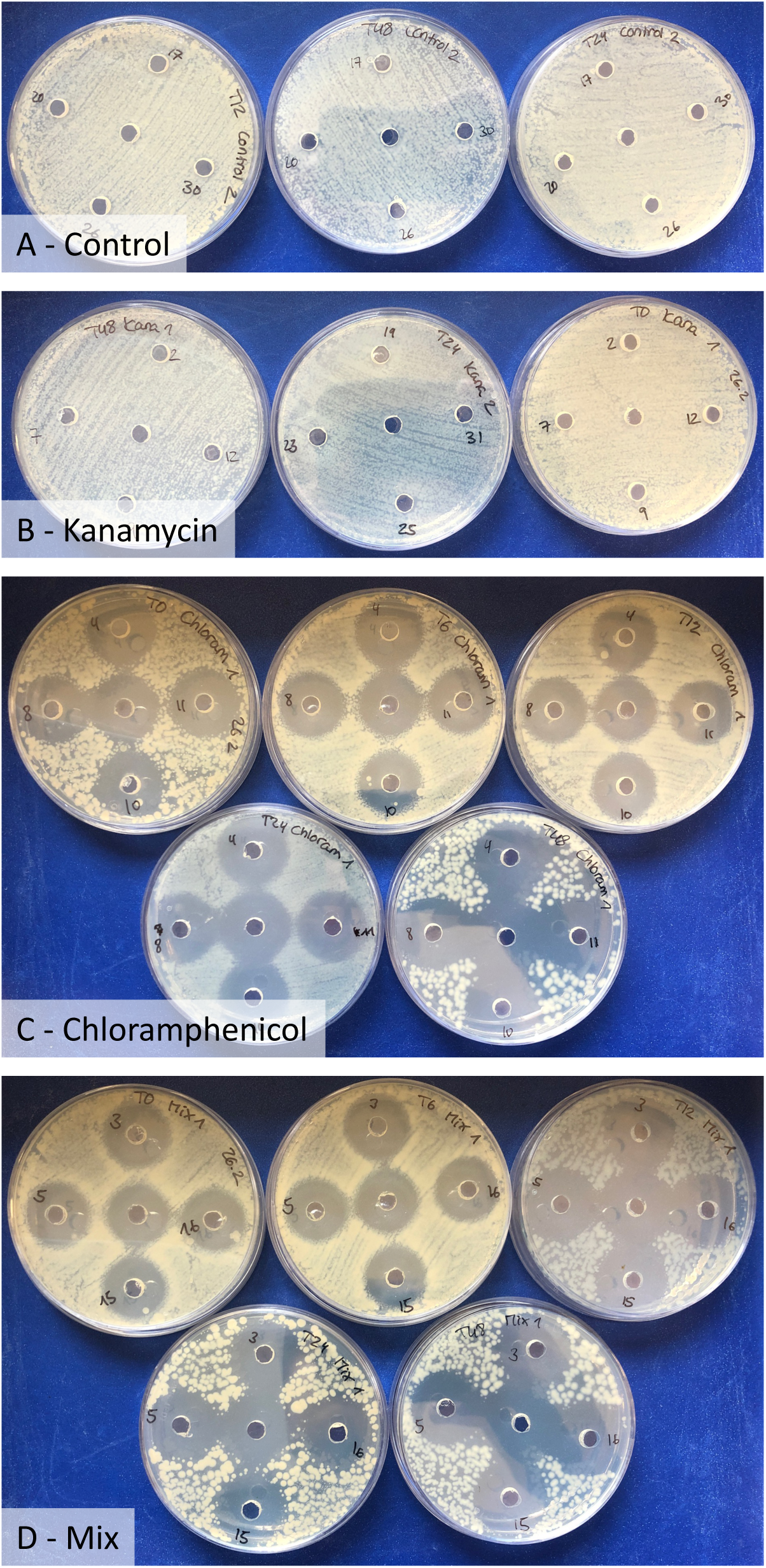
Well diffusion test to test for antibiotics in the water. Each well number corresponds to its respective tank. The center well was used for a control sample. Pictures were taken after 12 hours of incubation. Control (A) and Kanamycin (B) show no zones of inhibition, whereas treatments containing Chloramphenicol (C and D) show clear zones. Pictures show plates from beginning of the experiment (T0), after 24 hours (T24) and 48 hours (T48). For C and D additionally time points T6 and T12.

**Supplementary Figure 4:**
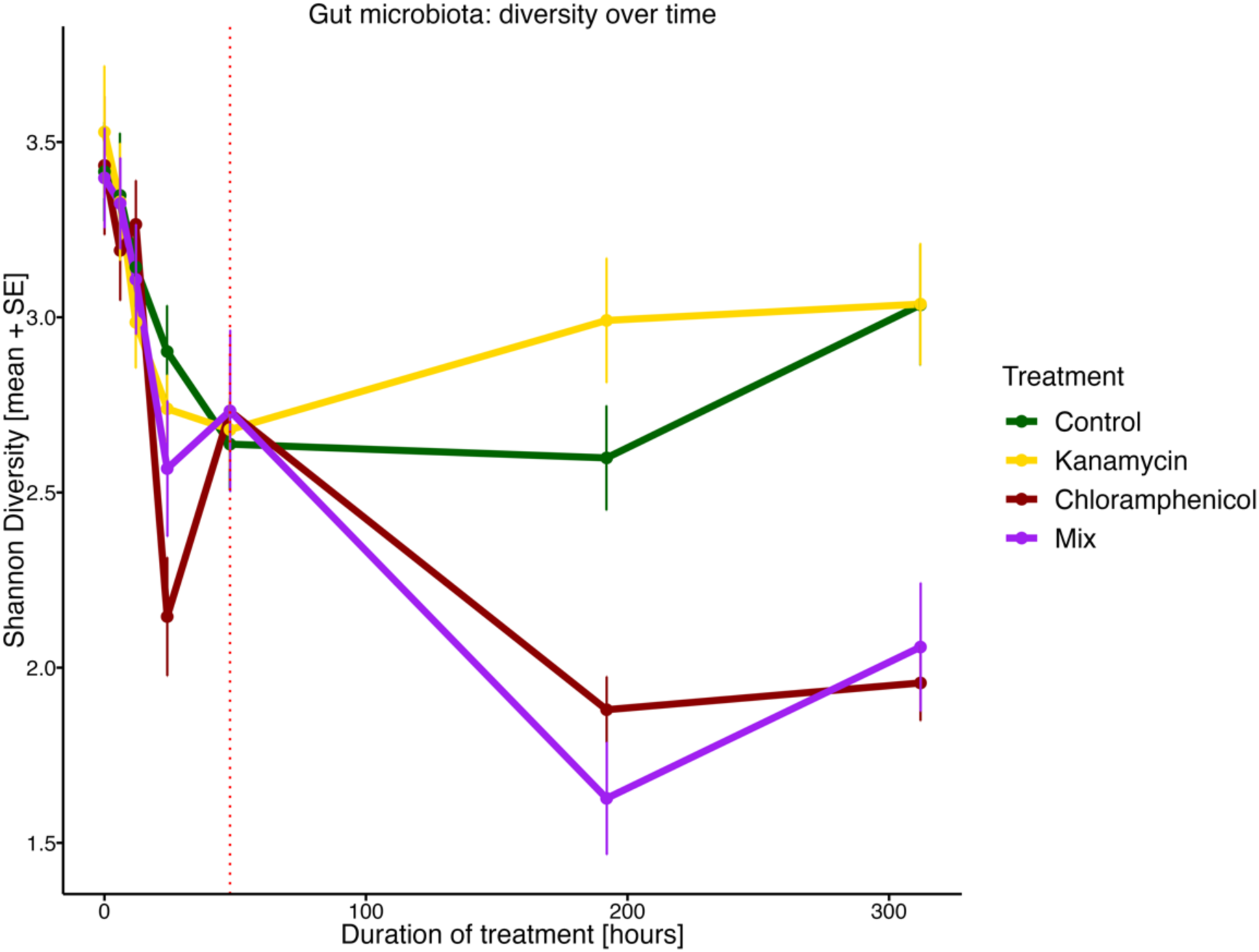
Alpha diversity changes over time under the influence of different antibiotic treatments in the gut microbiota of S. typhle. Colors represent treatment groups and data points display mean diversity ± standard error.

**Supplementary Figure 5:**
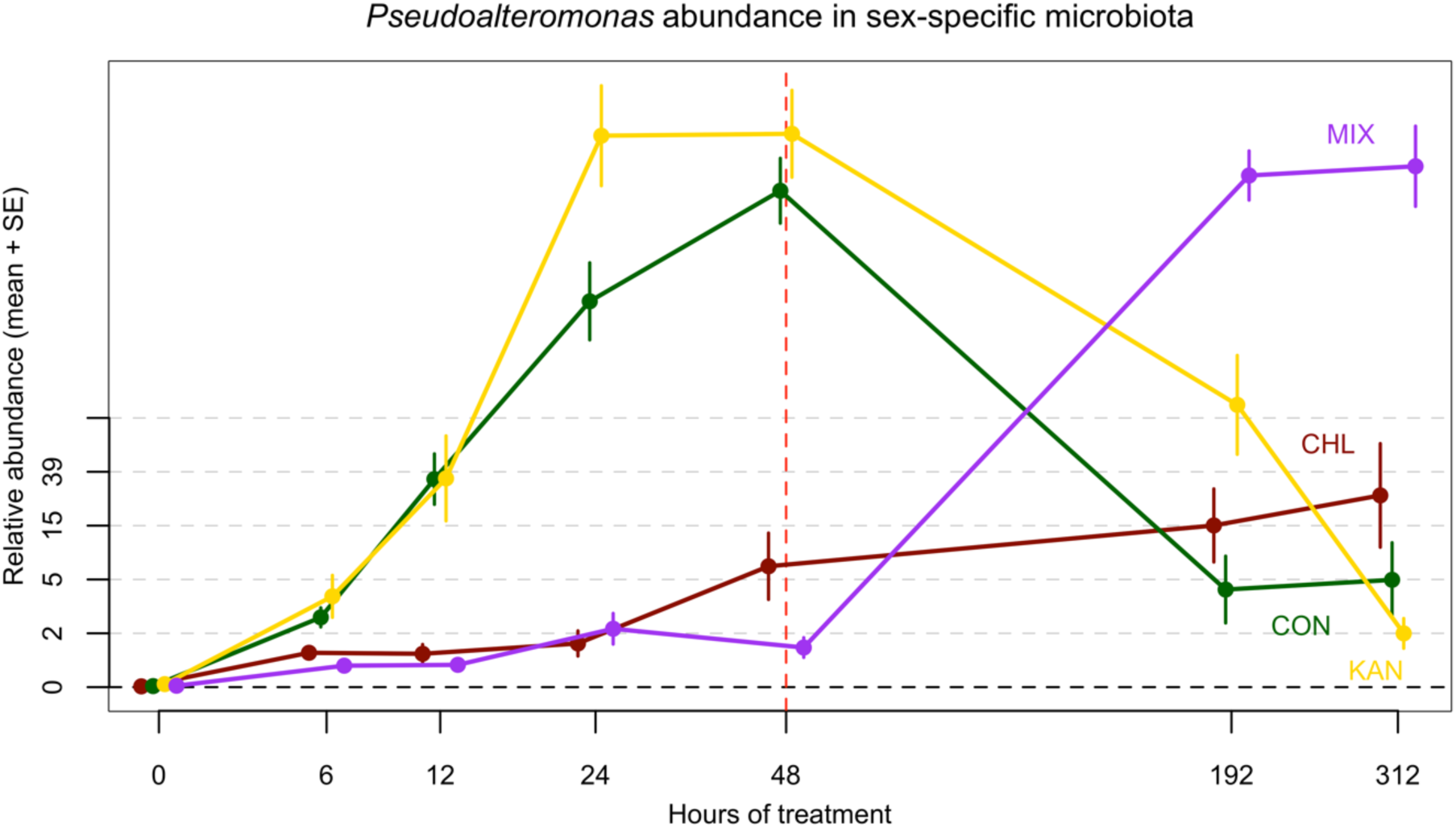
Pseudoalteromonas was initally selected as a paternal indicator species within the spike community but could not be depleted by the selected antibiotics and therefore excluded for further experiments.

**Supplementary Table 3:** Repeated measures PERMANOVA reveals significant impact of time, treatment and individual on microbial beta-diversity of the sex-specific microbiota.

|  | Df | SumOfSqs | MeanSqs | F.Model | R2 | Pr(>F) |
| --- | --- | --- | --- | --- | --- | --- |
| Treatment | 3 | 7.885 | 2.6282 | 13.308 | 0.06948 | 0.001 *** |
| Time | 1 | 7.962 | 7.9623 | 40.317 | 0.07016 | 0.001 *** |
| Individual | 52 | 33.077 | 0.6361 | 3.221 | 0.29146 | 0.001 *** |
| Treatment:Time | 3 | 8.87 | 2.9568 | 14.972 | 0.07816 | 0.001 *** |
| Residuals | 282 | 55.692 | 0.1975 |  | 0.49074 |  |
| Total | 341 | 113.486 |  |  | 1 |  |

**Supplementary Figure 6:**
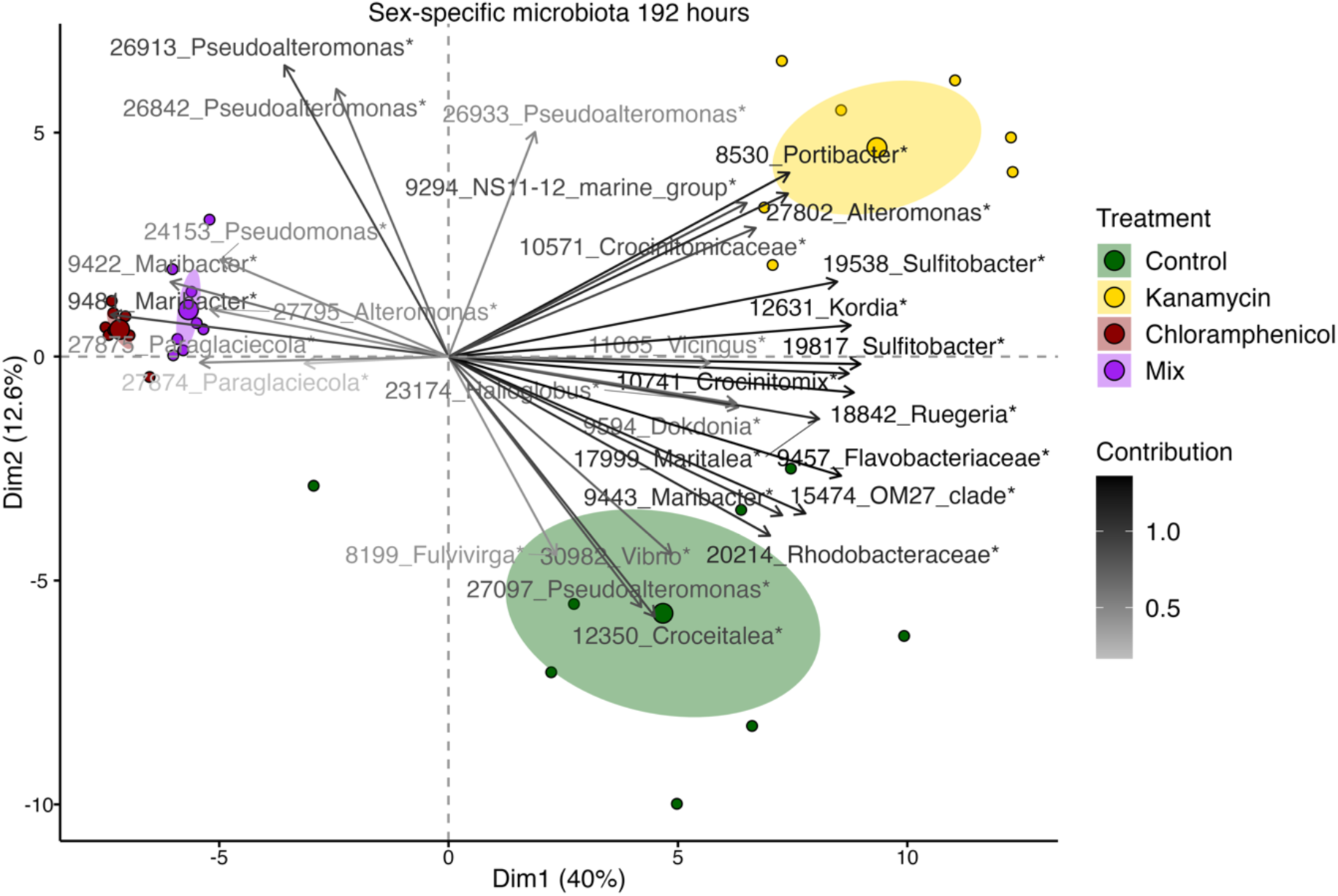
PCA Biplot of the sex-specific microbiota under the influence of different antibiotics at 192 hours of treatment. Ellipses display a 95% confidence interval around the group’s mean. Colors represent treatment groups. Factor map displays the ASV ID and the genus name, respectively family name if genus could not be identified.

<u>Experiment 3:</u> Manipulating paternal sex-specific microbiota to unravel impact on the offspring

**Supplementary Figure 7:**
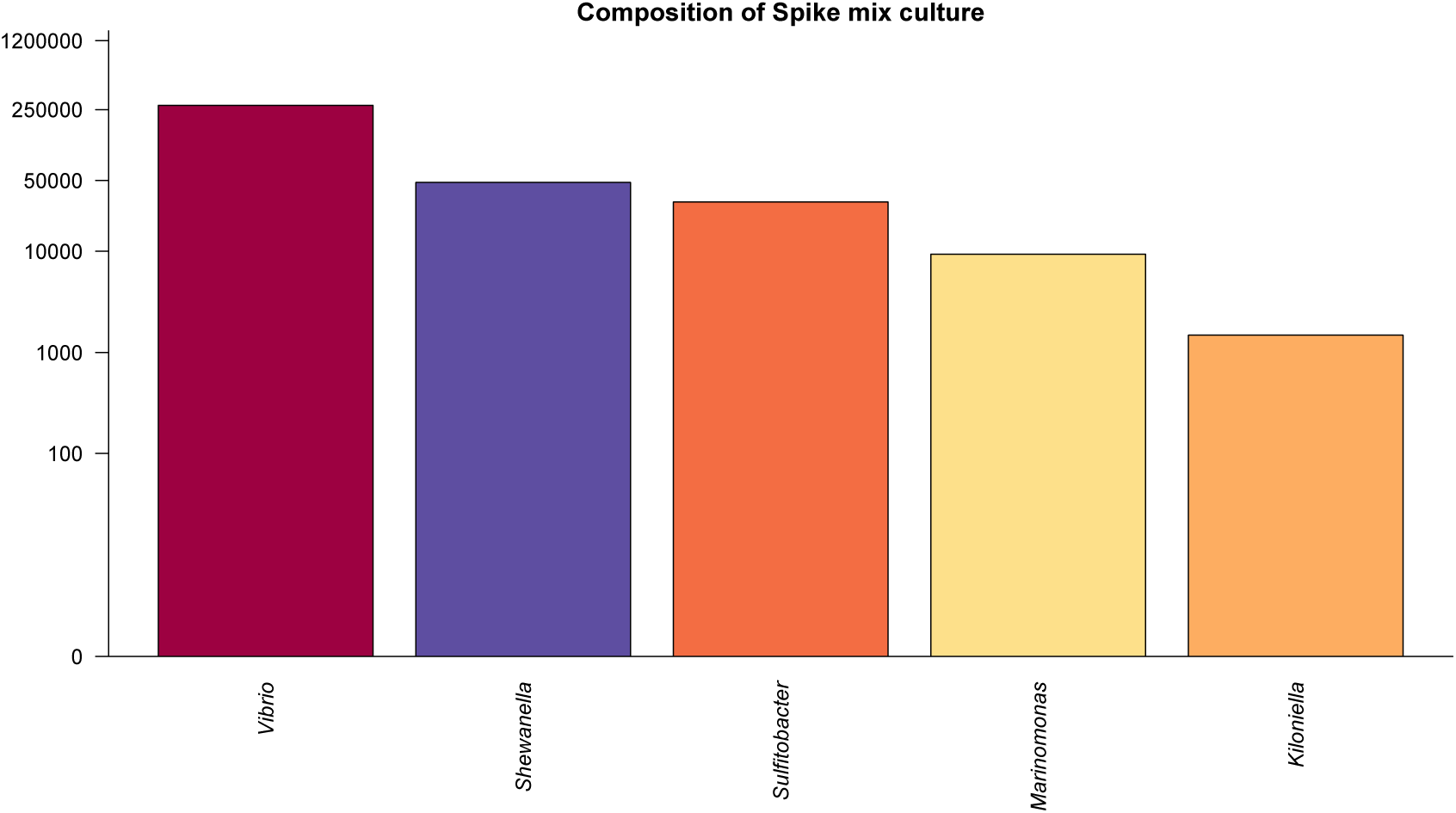
Composition of spike community mixed culture that was used in 30x concentration for treating the male brood pouch. Bars represent sequencing counts of selected candidate strains within the culture.

Experiment 3 (A) – Adults

**Supplementary Table 4:** Pregnancy statistic for all treatment groups. Spike_Treat includes fish of the treatment groups Spike and AntiSpike and Anti_Treat includes fish of the groups Antibiotics and AntiSpike.

| Treatment | Filling of brood pouch<br>(% + SD) | Gestation length<br>(mean days + SD) | Number of offspring<br>(mean + SD) |
| --- | --- | --- | --- |
| All fish | 86.76 ± 18.7 | 29.18 ± 1.62 | 31.76 ± 15.06 |
| Spike_Treat | 85.3 ± 21.76 | 28.47 ± 0.8 | 28.88 ± 14.69 |
| Anti_Treat | 90.63 ± 15.48 | 29.38 ± 1.63 | 33.63 ± 14.41 |
| Control | 91.67 ± 12.5 | 29.78 ± 1.86 | 36.11 ± 19.77 |
| Spike | 75 ± 25 | 28.22 ± 0.97 | 24.11 ± 7.85 |
| Antibiotics | 84.38 ± 18.6 | 30 ± 2.14 | 33 ± 9.12 |
| AntiSpike | 96.86 ± 8.84 | 28.75 ± 0.46 | 34.25 ± 18.99 |

**Supplementary Figure 8:**
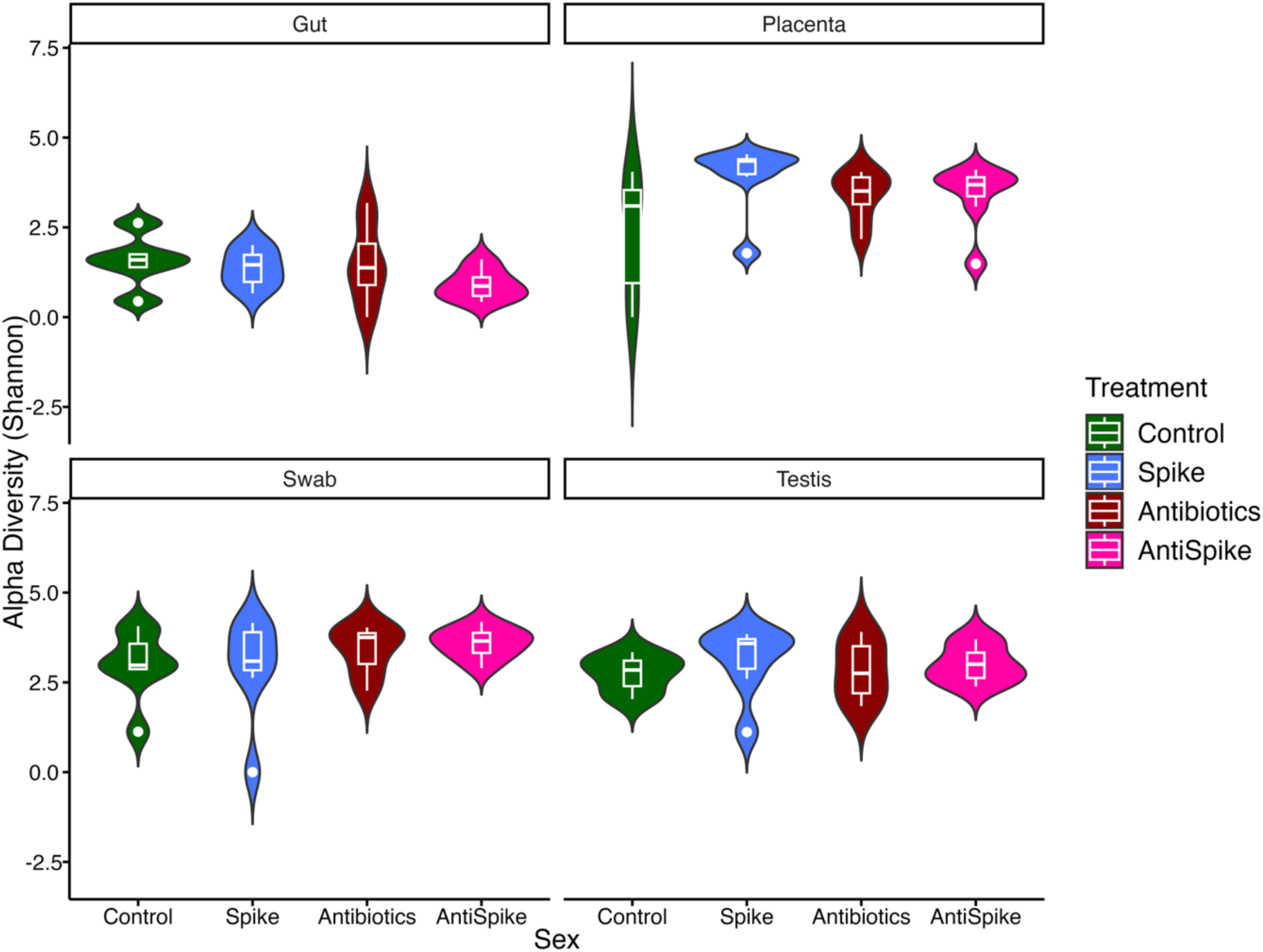
Violin and boxplots showing alpha diversity of male organs under the influence of different treatments. Panels indicate from which organ the microbiota was ribotyped.

**Supplementary Table 5:**
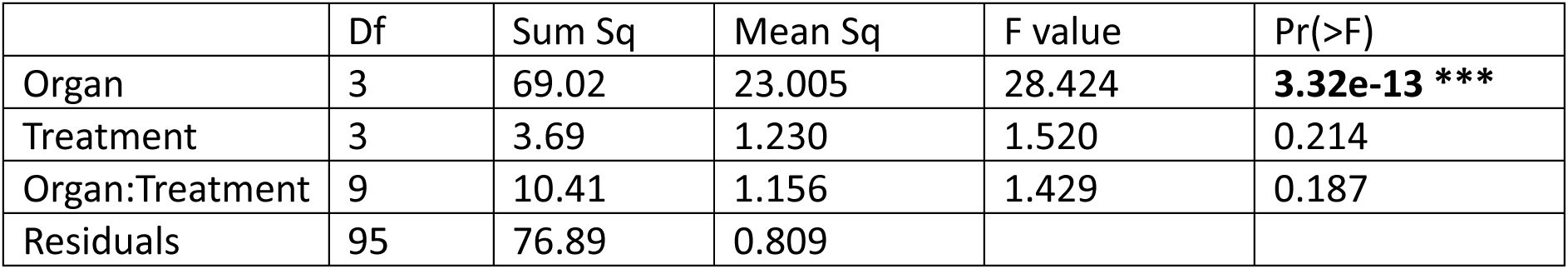
Two-way ANOVA comparing male organs under the influence of different treatments.

**Supplementary Figure 9:**
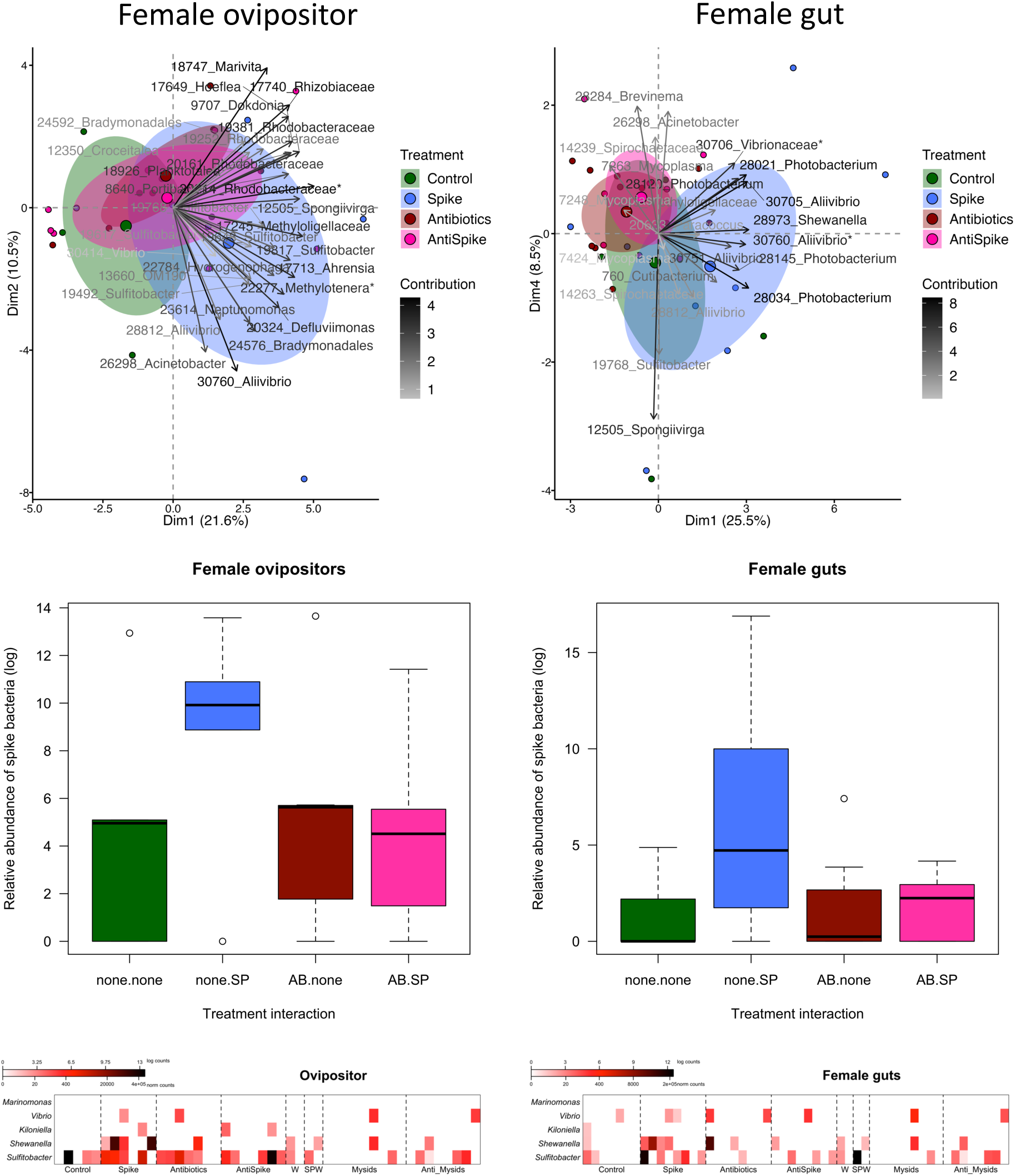
Female microbiome composition of ovipositor and gut, relative abundance of spike bacteria and counts of individual spike strains

**Supplementary Figure 10:**
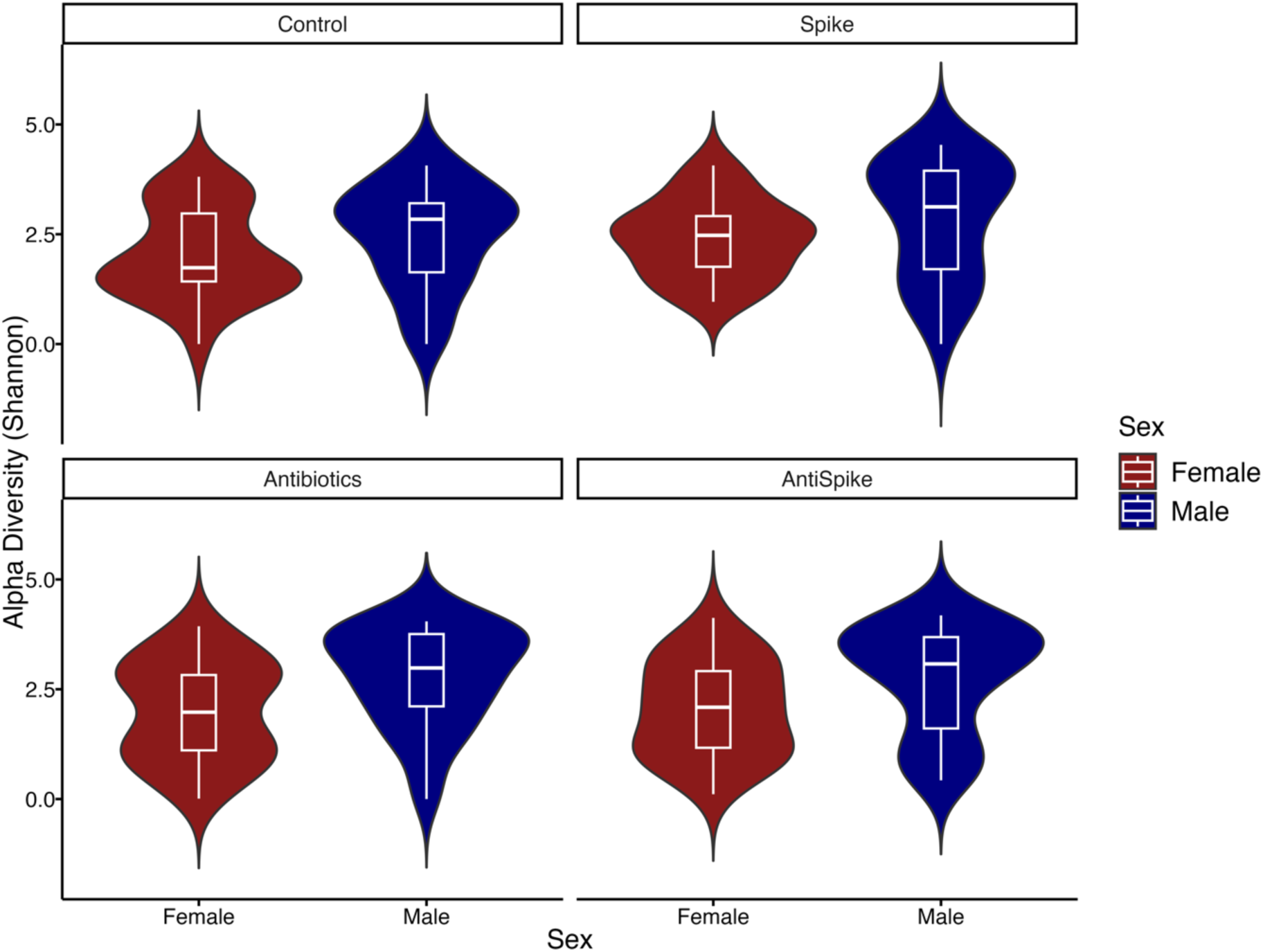
Violin and boxplot showing an alpha diversity comparison of females (red) and males (blue) separated by treatment group.

**Supplementary Figure 11:**
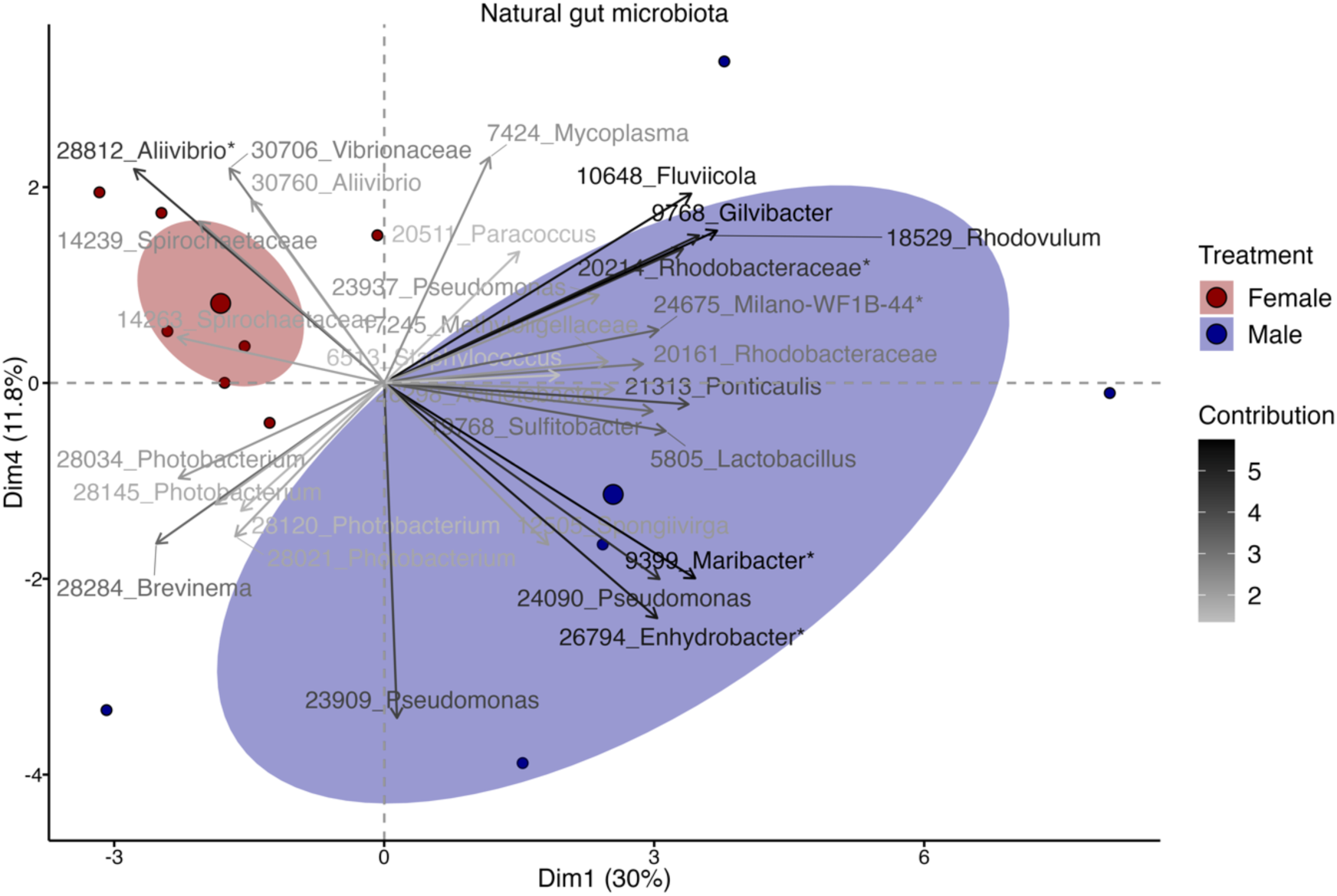
The natural gut microbiota of male and female pipefish (Control group only) indicates a strong sexual dimorphism. Ellipses indicate a confidence interval around the group’s mean. Factor map shows ASV ID and genus.

**Supplementary Table 6:** Two-way ANOVA results suggest a sexual dimorphism within the alpha diversity of male and female pipefish.

|  | Df | Sum Sq | Mean Sq | F value | Pr(>F) |
| --- | --- | --- | --- | --- | --- |
| Sex | 1 | 18.76 | 18.761 | 14.539 | **0.000181 \*\*\*** |
| Treatment | 3 | 3.43 | 1.142 | 0.885 | 0.449662 |
| Sex:Treatment | 3 | 1.45 | 0.485 | 0.376 | 0.770559 |
| Residuals | 206 | 265.82 | 1.290 |  |  |

**Supplementary Table 7:** One-way ANOVA results suggest a sexual dimorphism in the gut microbiota of pipefish (beta-diversity). Here, only the gut tissue of untreated males and females (Control group) was analyzed.

|  | Df | Pillai | Approx. F | Num Df | Den Df | Pr(>F) |
| --- | --- | --- | --- | --- | --- | --- |
| (Intercept) | 1 | 0.00000 | 0.00000 | 6 | 5 | 1.00000 |
| Sex | 1 | 0.89181 | 6.8689 | 6 | 5 | **0.02583 \*** |
| Residuals | 10 |  |  |  |  |  |

Experiment 3 (B) – Juveniles

**Supplementary Table 8:** Juvenile weight development is affected by time but not by treatment.

|  | Df | Sum Sq | Mean Sq | F value | Pr(>F) |
| --- | --- | --- | --- | --- | --- |
| Treatment | 3 | 54 | 18 | 0.333 | 0.802 |
| Time | 1 | 16961 | 16961 | 313.608 | **<2e-16 \*\*\*** |
| Treatment:Time | 3 | 238 | 79 | 1.466 | 0.242 |
| Residuals | 32 | 1731 | 54 |  |  |

**Supplementary Table 9:** Juvenile size development is strongly affected by time and time*treatment interaction but not by treatment alone.

|  | Df | Sum Sq | Mean Sq | F value | Pr(>F) |
| --- | --- | --- | --- | --- | --- |
| Treatment | 3 | 7 | 2 | 1.150 | 0.3441 |
| Time | 1 | 4968 | 4968 | 2380.310 | **<2e-16 \*\*\*** |
| Treatment:Time | 3 | 19 | 6 | 2.991 | **0.0454 \*** |
| Residuals | 32 | 67 | 2 |  |  |

**Supplementary Table 10:**
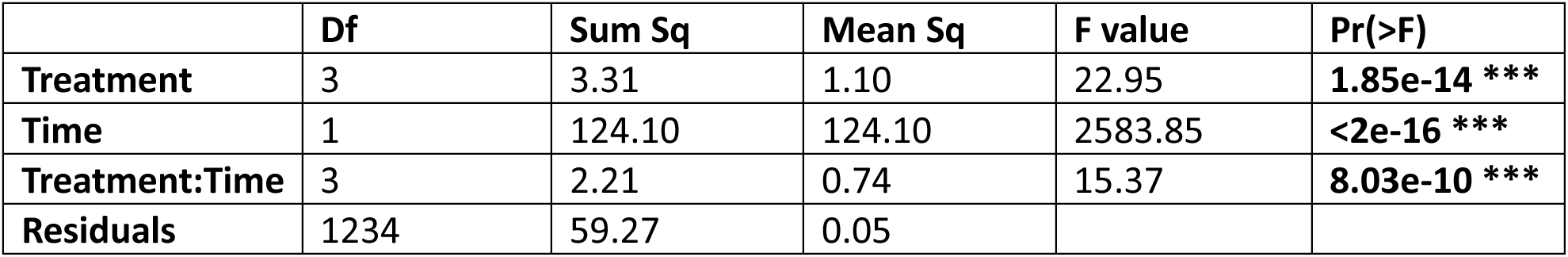
Juvenile survival is strongly affected by treatment, time and their interaction.

**Supplementary Table 11:** Contrasts (pairwise, adjusted by TukeyHSD) reveal that offspring of spike-treated fathers have a significantly higher survival.

| Contrast | Estimate | SE | df | t.ratio | p.value |
| --- | --- | --- | --- | --- | --- |
| Antibiotics-AntiSpike | 0.0230 | 0.0184 | 1234 | 1.251 | 0.5945 |
| Antibiotics-Control | 0.0451 | 0.0162 | 1234 | 2.792 | 0.0273 * |
| Antibiotics-Spike | -0.0942 | 0.0175 | 1234 | -5.398 | <.0001 *** |
| AntiSpike-Control | 0.0221 | 0.0184 | 1234 | 1.198 | 0.6283 |
| AntiSpike-Spike | -0.1172 | 0.0196 | 1234 | -5.992 | <.0001 *** |
| Control-Spike | -0.1393 | 0.0175 | 1234 | -7.983 | <.0001 *** |

**Supplementary Table 12:** Whole-body microbiota: ANCOVA testing the influence of treatment and/or time on microbiota composition (alpha-diversity).

|  | Estimate | Std. Error | t value | Pr(>\|t\|) |
| --- | --- | --- | --- | --- |
| (Intercept) | 2.878693 | 0.128019 | 22.486 | <2e-16 *** |
| AntiSpike | 0.026886 | 0.163318 | 0.165 | 0.8694 |
| Control | -0.335738 | 0.161476 | -2.079 | **0.0386 \*** |
| Spike | -0.060166 | 0.164025 | -0.367 | 0.7141 |
| TimePoint | 0.002756 | 0.004850 | 0.568 | 0.5704 |

**Supplementary Table 13:** Gut microbiota: ANCOVA testing the influence of treatment and/or time on microbiota composition (alpha-diversity).

|  | Estimate | Std. Error | t value | Pr(>\|t\|) |
| --- | --- | --- | --- | --- |
| (Intercept) | 3.092842 | 0.133810 | 23.114 | < 2e-16 *** |
| AntiSpike | 0.127106 | 0.170633 | 0.745 | 0.457 |
| Control | -0.269765 | 0.169415 | -1.592 | 0.113 |
| Spike | -0.246375 | 0.171374 | -1.438 | 0.152 |
| TimePoint | -0.022126 | 0.004995 | -4.430 | **1.4e-05 \*\*\*** |

**Supplementary Table 14:** Beta-diversity: ANOVA p-values evaluating the MANOVA model on the principal components (PCs) of the external whole-body microbiome of juveniles at each time point. Comparison of Antibiotics (AB) treatment groups (Antibiotics and AntiSpike) and Spike (SP) treatment groups (Spike and AntiSpike).

|  |  |  |  |  |  |  |  |  |  |  |
| --- | --- | --- | --- | --- | --- | --- | --- | --- | --- | --- |
| dpr | 0 | 2 | 4 | 6 | 8 | 10 | 12 | 14 | 30 | 45 |
| AB | 0.386 | 0.052 | **0.043** | **0.039** | 0.636 | **0.025** | 0.094 | 0.717 | **0.025** | 0.501 |
| SP | 0.204 | **0.013** | **0.035** | **0.006** | 0.302 | **0.017** | 0.299 | **0.015** | **0.009** | 0.308 |
| AB:SP | 0.826 | 0.943 | 0.261 | 0.267 | 0.602 | 0.883 | 0.084 | **0.039** | 0.251 | 0.411 |

**Supplementary Figure 12:**
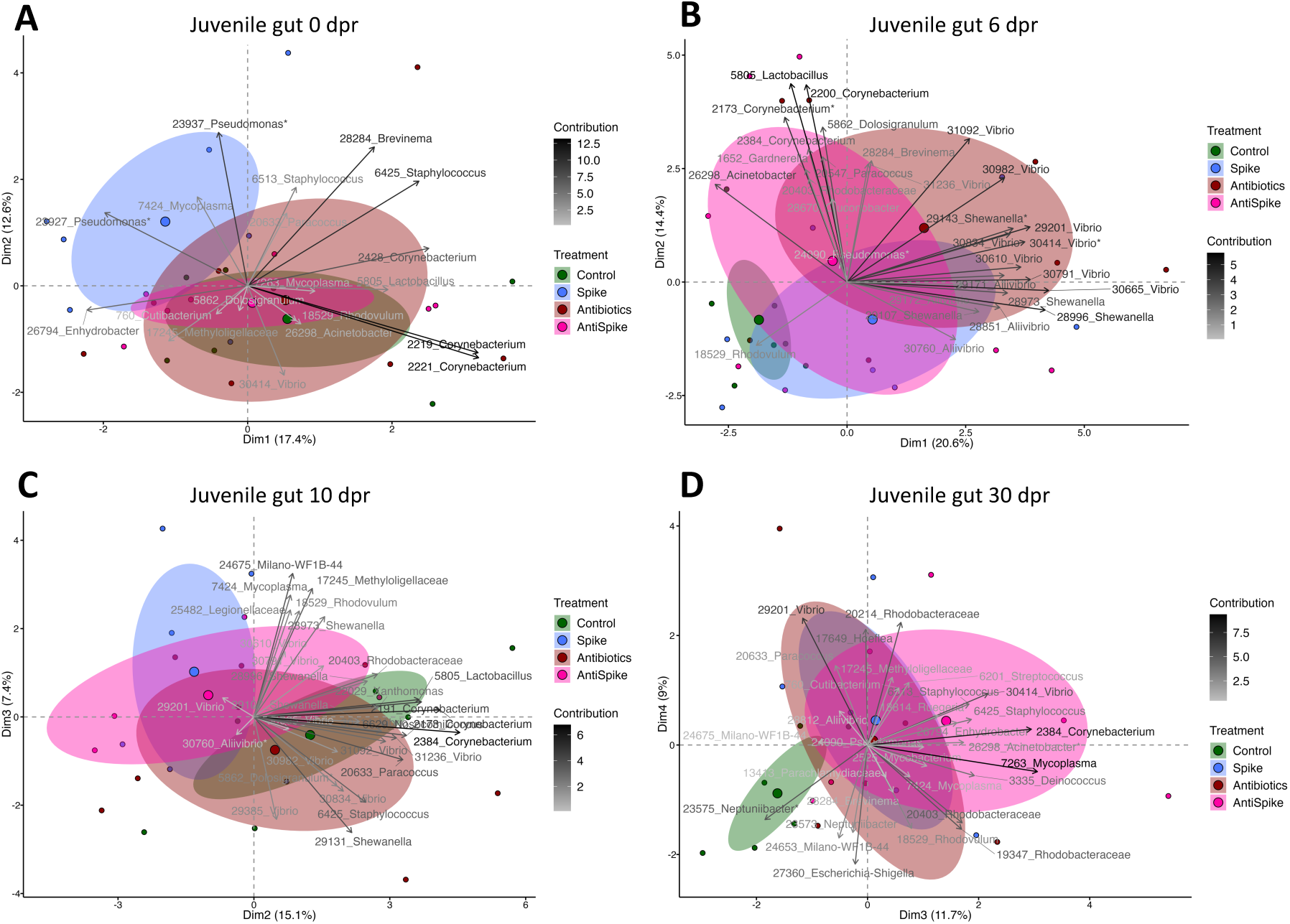
Internal (gut) microbiota composition of juveniles from birth (A), 6 days (B), 10 days (C) up to 30 days (D) after birth. Colors represent paternal treatment. Ellipses show a 95% confidence interval around the group’s mean. Factor map shows ASV ID and genus name respectively family name if genus could not be identified.

**Supplementary Table 15:** Beta diversity: ANOVA p-values evaluating the MANOVA model on the principal components (PCs) of the internal sterile gut microbiome of juveniles at each time point. Comparison of Antibiotics (AB) treatment groups (Antibiotics and AntiSpike) and Spike (SP) treatment groups (Spike and AntiSpike).

|  |  |  |  |  |  |  |  |  |  |  |
| --- | --- | --- | --- | --- | --- | --- | --- | --- | --- | --- |
| dpr | 0 | 2 | 4 | 6 | 8 | 10 | 12 | 14 | 30 | 45 |
| AB | 0.646 | 0.399 | 0.435 | 0.185 | 0.245 | 0.875 | 0.135 | 0.923 | 0.332 | 0.218 |
| SP | 0.325 | 0.467 | 0.488 | 0.941 | 0.591 | 0.291 | **0.044** | 0.924 | 0.366 | 0.558 |
| AB:SP | 0.471 | 0.290 | 0.244 | 0.336 | 0.422 | 0.975 | 0.481 | 0.348 | 0.609 | 0.707 |

**Experiment 3 (C) – Spike community**

**Supplementary Figure 13:**
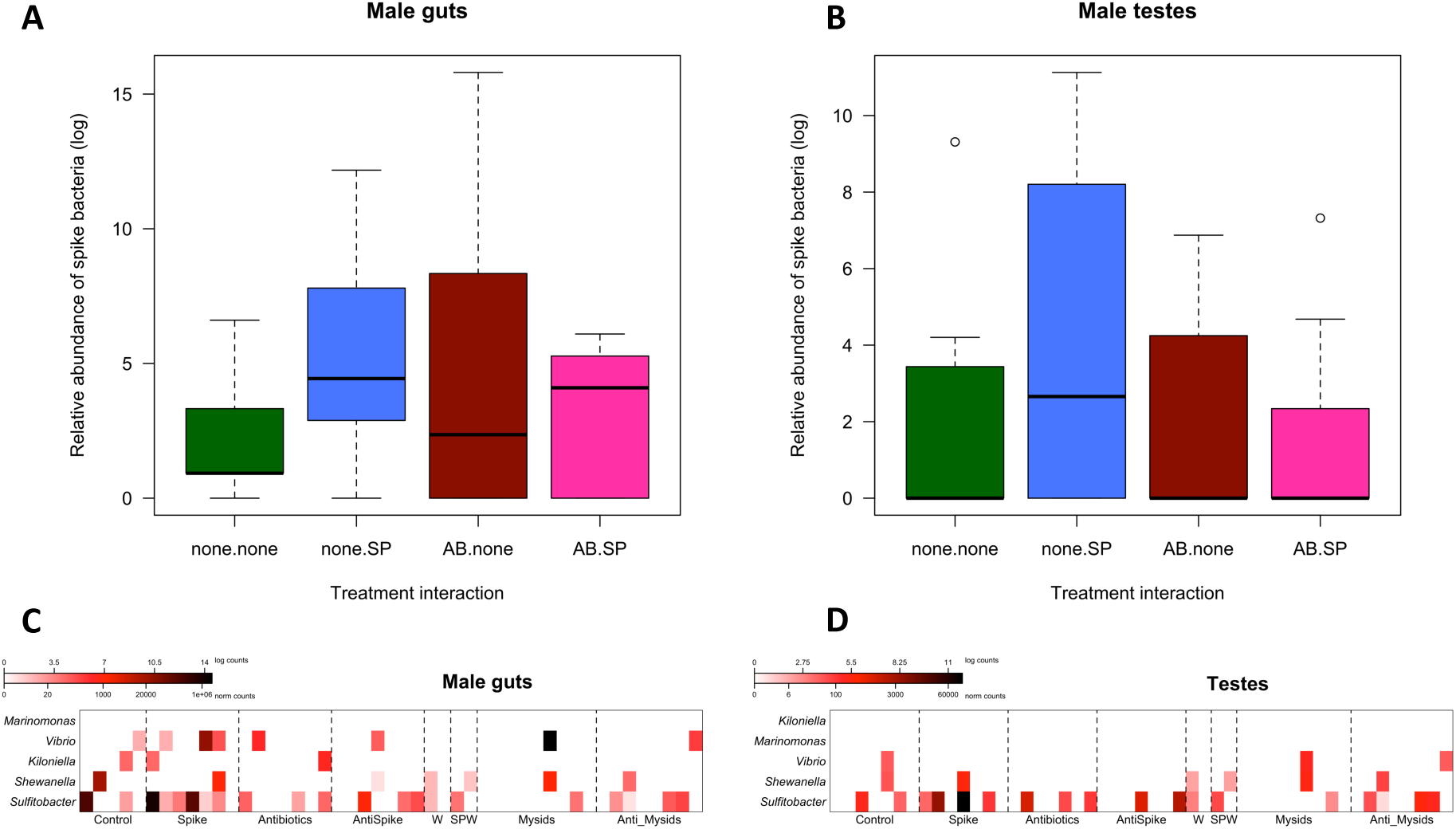
Relative abundance of spike community strains in the male gut (A) and the male testes (B). Heatmaps showing counts of individual strains for gut (C) and testes (D).

**Supplementary Table 16:** Kruskal-Wallis H test p-values indicate impact of spike community strains on adult tissues of S. typhle. Data is based on normalized counts.

|  | Placenta-like<br>tissue | Brood pouch<br>swabs | Male gut | Male<br>testes | Ovipositor | Female gut |
| --- | --- | --- | --- | --- | --- | --- |
| All strains | **0.021** | **0.044** | 0.217 | 0.637 | 0.250 | 0.080 |
| *Kiloniella* | 0.433 | 0.321 | 0.532 | *n/a* | 0.368 | 0.392 |
| *Marinomonas* | *n/a* | 0.392 | *n/a* | *n/a* | *n/a* | *n/a* |
| *Sulfitobacter* | 0.091 | 0.760 | 0.817 | 0.542 | 0.286 | 0.082 |
| *Vibrio* | 0.258 | 0.240 | 0.219 | 0.392 | 0.392 | 0.708 |
| *Shewanella* | 0.392 | **0.018** | 0.392 | 0.392 | 0.179 | 0.430 |

## Notes

### Competing Interest Statement

The authors have declared no competing interest.

## References

Altschul, S. F., T. L. Madden, A. A. Schäffer, J. Zhang, Z. Zhang, W. Miller, and D. J. Lipman. 1997. Gapped BLAST and PSI-BLAST: A new generation of protein database search programs. Nucleic Acids Res 25:3389–3402.

Balouiri, M., M. Sadiki, and S. K. Ibnsouda. 2016. Methods for in vitro evaluating antimicrobial activity: A review. J Pharm Anal 6:71–79.

Beemelmanns, A., M. Poirier, T. Bayer, S. Kuenzel, and O. Roth. 2019. Microbial embryonal colonization during pipefish male pregnancy. Sci Rep 9:1–14. Springer US.

Beemelmanns, A., and O. Roth. 2016. Bacteria-type-specific biparental immune priming in the pipefish Syngnathus typhle. Ecol Evol 6:6735–6757.

Beemelmanns, A., and O. Roth. 2017. Grandparental immune priming in the pipefish Syngnathus typhle Grandparental immune priming in the pipefish Syngnathus typhle. BMC Evol Biol, doi: 10.1186/s12862-017-0885-3. BMC Evolutionary Biology.

Blackburn, D. G. 2015. Evolution of vertebrate viviparity and specializations for fetal nutrition: A quantitative and qualitative analysis. J Morphol 276:961–990.

Bogaert, D., G. J. van Beveren, E. M. de Koff, P. Lusarreta Parga, C. E. Balcazar Lopez, L. Koppensteiner, M. Clerc, R. Hasrat, K. Arp, M. L. J. N. Chu, P. C. M. de Groot, E. A. M. Sanders, M. A. van Houten, and W. A. A. de Steenhuijsen Piters. 2023. Mother-to-infant microbiota transmission and infant microbiota development across multiple body sites. Cell Host Microbe 31:447-460.e6. Cell Press.

Bouchet, V., H. Huot, and R. Goldstein. 2008. Molecular Genetic Basis of Ribotyping. Clin Microbiol Rev 21:262–273.

Cáceres, M. De, and P. Legendre. 2009. Associations between species and groups of sites: indices and statistical inference. Ecology 90:3566–3574.

Callahan, B. J., P. J. McMurdie, M. J. Rosen, A. W. Han, A. J. A. Johnson, and S. P. Holmes. 2016. DADA2: High-resolution sample inference from Illumina amplicon data. Nat Methods 13:581–583.

Daskalakis, G., A. Psarris, A. Koutras, Z. Fasoulakis, I. Prokopakis, A. VarthaliP, C. Karasmani, T. Ntounis, E. Domali, M. Theodora, P. Antsaklis, K. I. Pappa, and A. Papapanagiotou. 2023. Maternal Infection and Preterm Birth: From Molecular Basis to Clinical Implications. Children 10:907.

de la Cuesta-Zuluaga, J., S. T. Kelley, Y. Chen, J. S. Escobar, N. T. Mueller, R. E. Ley, D. McDonald, S. Huang, A. D. Swafford, R. Knight, and V. G. Thackray. 2019. Age-and Sex-Dependent Paierns of Gut Microbial Diversity in Human Adults. mSystems 4.

Donald, K., and B. B. Finlay. 2023. Early-life interactions between the microbiota and immune system: impact on immune system development and atopic disease. Nat Rev Immunol 23:735–748.

He, J., J. Lange, G. Marinos, J. Bathia, D. Harris, R. Soluch, V. Vaibhvi, P. Deines, M. A. Hassani, K.-S. Wagner, R. Zapien-Campos, C. Jaspers, and F. Sommer. 2020. Advancing Our Functional Understanding of Host-Microbiota Interactions: A Need for New Types of Studies. BioEssays 1900211:1900211.

Lê, S., J. Josse, and F. Husson. 2008. FactoMineR: An R package for multivariate analysis. J Stat Sokw 25:1–18.

Lemaire, O. N., V. Méjean, and C. Iobbi-Nivol. 2020. The Shewanella genus: ubiquitous organisms sustaining and preserving aquatic ecosystems. FEMS Microbiol Rev 44:155–170.

McMurdie, P. J., and S. Holmes. 2013. phyloseq: An R Package for Reproducible Interactive Analysis and Graphics of Microbiome Census Data. PLoS One 8:e61217.

Moia, E. V. S., and N. A. Moran. 2024. The honeybee microbiota and its impact on health and disease. Nat Rev Microbiol 22:122–137.

Munro, P. D., A. Barbour, and T. H. Birkbeck. 1995. Comparison of the Growth and Survival of Larval Turbot in the Absence of Culturable Bacteria with Those in the Presence of Vibrio anguillarum, Vibrio alginolyticus, or a Marine Aeromonas sp. Appl Environ Microbiol 61:4425–4428.

Nyholm, S. V. 2020. In the beginning: egg–microbe interactions and consequences for animal hosts. Philosophical Transactions of the Royal Society B: Biological Sciences 375:20190593.

Pappert, F. A., A. Dubin, G. G. Torres, and O. Roth. 2024. Navigating sex and sex roles: deciphering sex-biased gene expression in a species with sex-role reversal (Syngnathus typhle). R Soc Open Sci 11.

Quast, C., E. Pruesse, P. Yilmaz, J. Gerken, T. Schweer, P. Yarza, J. Peplies, and F. O. Glöckner. 2013. The SILVA ribosomal RNA gene database project: Improved data processing and web-based tools. Nucleic Acids Res 41:590–596.

R Core Team. 2024. R: A language and environment for statistical computing.

Roth, O., V. Klein, A. Beemelmanns, J. P. Scharsack, and T. B. H. Reusch. 2012. Male pregnancy and biparental immune priming. American Naturalist 180:802–814.

Stölting, K. N., and A. B. Wilson. 2007. Male pregnancy in seahorses and pipefish: Beyond the mammalian model. BioEssays 29:884–896.

Tanger, I. S., J. Stefanschitz, Y. Schwert, and O. Roth. 2024. The source of microbial transmission influences niche colonization and microbiome development. Proceedings of the Royal Society B: Biological Sciences 291. Royal Society Publishing.

Valeri, F., and K. Endres. 2021. How biological sex of the host shapes its gut microbiota. Front Neuroendocrinol 61:100912.

Verner-Jeffreys, D. W., R. J. Shields, and T. H. Birkbeck. 2003. Bacterial influences on Atlantic halibut Hippoglossus hippoglossus yolk-sac larval survival and start-feed response. Dis Aquat Organ 56:105–113.

Větrovský, T., and P. Baldrian. 2013. The Variability of the 16S rRNA Gene in Bacterial Genomes and Its Consequences for Bacterial Community Analyses. PLoS One 8:e57923.

Vitikainen, E. I. K., M. Meniri, H. H. Marshall, F. J. Thompson, R. Businge, F. Mwanguhya, S. Kyabulima, K. Mwesige, S. Ahabonya, J. L. Sanderson, G. Kalema-Zikusoka, J. I. Hoffman, D. Wells, G. Lewis, S. L. Walker, H. J. Nichols, J. D. Blount, and M. A. Cant. 2023. The social formation of fitness: lifetime consequences of prenatal nutrition and postnatal care in a wild mammal population. Philosophical Transactions of the Royal Society B: Biological Sciences 378.

Wendling, C. C., A. Piecyk, D. Refardt, C. Chibani, R. Hertel, H. Liesegang, B. Bunk, J. Overmann, and O. Roth. 2017. Tripartite species interaction: eukaryotic hosts suffer more from phage susceptible than from phage resistant bacteria. BMC Evol Biol 17:1–12. BMC Evolutionary Biology.

Whitngton, C. M., and C. R. Friesen. 2020. The evolution and physiology of male pregnancy in syngnathid fishes. Biological Reviews 95:1252–1272. Blackwell Publishing Ltd.

Wosnick, N., R. D. Leite, E. P. Giareta, D. Morick, and R. A. Hauser-Davis. 2022. Unraveling Metabolite Provisioning to Offspring Through Parental Fluids: A Case Study of the Brazilian Guitarfish, Pseudobatos horkelii. Front Physiol 13.

